# Genetic regulation of circulating metabolome in cattle

**DOI:** 10.64898/2026.09.05.749592

**Authors:** Jun Teng, Houcheng Li, Jian Yang, Jingsheng Lu, Chongwei Duan, Zhujun Chen, Xinyi Zhang, Xiuxin Zhao, Fen Pei, Xiaoping Wu, Pengju Zhao, Haihan Zhang, Hongding Gao, Lewei Guo, Dan Wang, Chao Ning, Huiming Liu, Guosheng Su, Rongling Li, Yundong Gao, Jianbin Li, Qin Zhang, Lingzhao Fang, Xiao Wang

## Abstract

The circulating metabolome is a vital intermediate layer linking genetics to complex phenotypes, yet existing genome-wide studies—even in humans—rarely capture dynamic physiological contexts or cellular regulatory mechanisms. Here, we present the **Cattle Metabolome Atlas** (https://cattlema.farmgtex.org/), a comprehensive resource of 3,436 plasma metabolites and 4,851 serum metabolites from 4,651 animals with matched sequence-level genotypes across highly dynamic parity and lactation stages. We mapped 728 plasma and 622 serum metabolite quantitative trait loci (mQTL), revealing widespread context-dependent regulatory architectures. Integrating these mQTL with the multi-tissue expression quantitative trait loci (eQTL) and single-cell atlas of 59 tissues demonstrates that 74–79% of mQTL colocalize with eQTL across 29 tissues, prioritizing the liver as the primary systemic hub and resolving metabolic programs at single-cell resolution. Furthermore, we charted 538 causal gene-tissue-metabolite-trait cascades across 11 complex traits in cattle. Finally, cross-species analyses demonstrate partial evolutionary conservation of genetic metabolism between cattle and humans, establishing this atlas as a powerful asset for cattle genetics and genomics, selective breeding, and comparative biology.

## Introduction

The metabolome represents the complete quantitative collection of small molecules within an organism, providing a direct, proximal biochemical readout of cellular activity, physiological states, and metabolic reactions^1^. Because metabolites lie closer to organismal phenotypes than upstream transcripts, disease-associated variants identified via genome-wide association studies (GWAS) are frequently enriched for metabolite quantitative trait loci (mQTL)^2^. Recent large-scale metabolite GWAS (mGWAS) have identified numerous mQTL across diverse classes of circulating molecules, positioning genetically regulated metabolites not merely as biomarkers, but as crucial mechanistic anchors for dissecting the molecular basis of complex traits^3–10^. While these foundational studies have primarily focused on common variants, emerging evidence indicates that rare variants also contribute substantially to metabolic variance^9,11–13^, suggesting that the full allelic spectrum must be explored to comprehensively resolve mammalian metabolic regulation.

Cattle represent an exceptional, yet underutilized, large-animal model for mammalian physiology and comparative medicine due to their strong physiological and immune-metabolic similarities to humans^14–18^. Driven by intensive artificial selection for production traits, dairy cattle undergo some of the most extreme metabolic adaptations observed in any mammalian species during the transition from late pregnancy to early lactation, a dynamic context characterized by a profound negative energy balance and systemic tissue remodeling that remains virtually unstudied at scale in other mammals^19,20^. Mapping these extreme metabolic states provides an invaluable resource for detecting mechanisms underlying milk production traits and other relevant metabolic disorders (e.g., ketosis and hypocalcemia). The findings and insights can also serve as evolutionarily conserved window into human clinical pathologies, offering therapeutic insights for female reproductive health disorders, inflammatory conditions, and metabolic syndrome^21–24^. However, our understanding of the genetic architecture governing these processes remains fragmented; previous livestock mGWAS have been severely limited by small cohort sizes, narrow metabolite coverage, a complete lack of context-specific and tissue/cell-type resolutions, and a historical absence of paired whole-genome sequencing (WGS) data required to capture the regulatory contributions of rare variants^25–30^.

To overcome these systemic limitations and map the context-specific genetic architecture of a large mammal metabolome, we present the Cattle Metabolome Atlas (CattleMA, **Fig.** 1), established as a core initiative of the Farm Animal Genotype-Tissue Expression (FarmGTEx) project^31^. We deployed an expansive study design comprising 4,651 blood samples (3,653 plasma and 998 serum profiles) tracking cohorts across highly dynamic parity and lactation contexts, paired with a multi-tier sequencing strategy, integrating SNP arrays (*n* = 1,741), low-coverage WGS (LCWGS, *n* = 1,475), and high-coverage WGS (HCWGS, *n* = 1,435), and high-resolution, untargeted liquid chromatography-mass spectrometry (LC-MS) metabolomics. By leveraging this multi-layered resource, we map common and context-specific mQTL across shifting physiological states, utilize gene-based collapsing analyses to uncover the regulatory influence of rare/low-frequency variants, and integrate these signals with multi-tissue eQTL from the Cattle Genotype-Tissue Expression (CattleGTEx)^32^ and single-cell atlas of 59 tissues from the Cattle Cell Atlas^33^ to explore the molecular mechanisms underlying these context-specific mQTL. By examining GWAS results of 11 complex traits of economic importance in cattle, we further explored gene-tissue-cell-metabolite regulatory circuitry underlying complex traits. Finally, comparative analyses between cattle and humans highlighted evolutionary conservation of genetic regulation of metabolites, supporting the utility of cattle as a potential model for understanding human metabolic traits and diseases. CattleMA, together with its publicly accessible database (https://cattlema.farmgtex.org/), provides a comprehensive and valuable resource for cattle genetics, functional genomics, precision breeding, and comparative biology.

**Fig 1.**
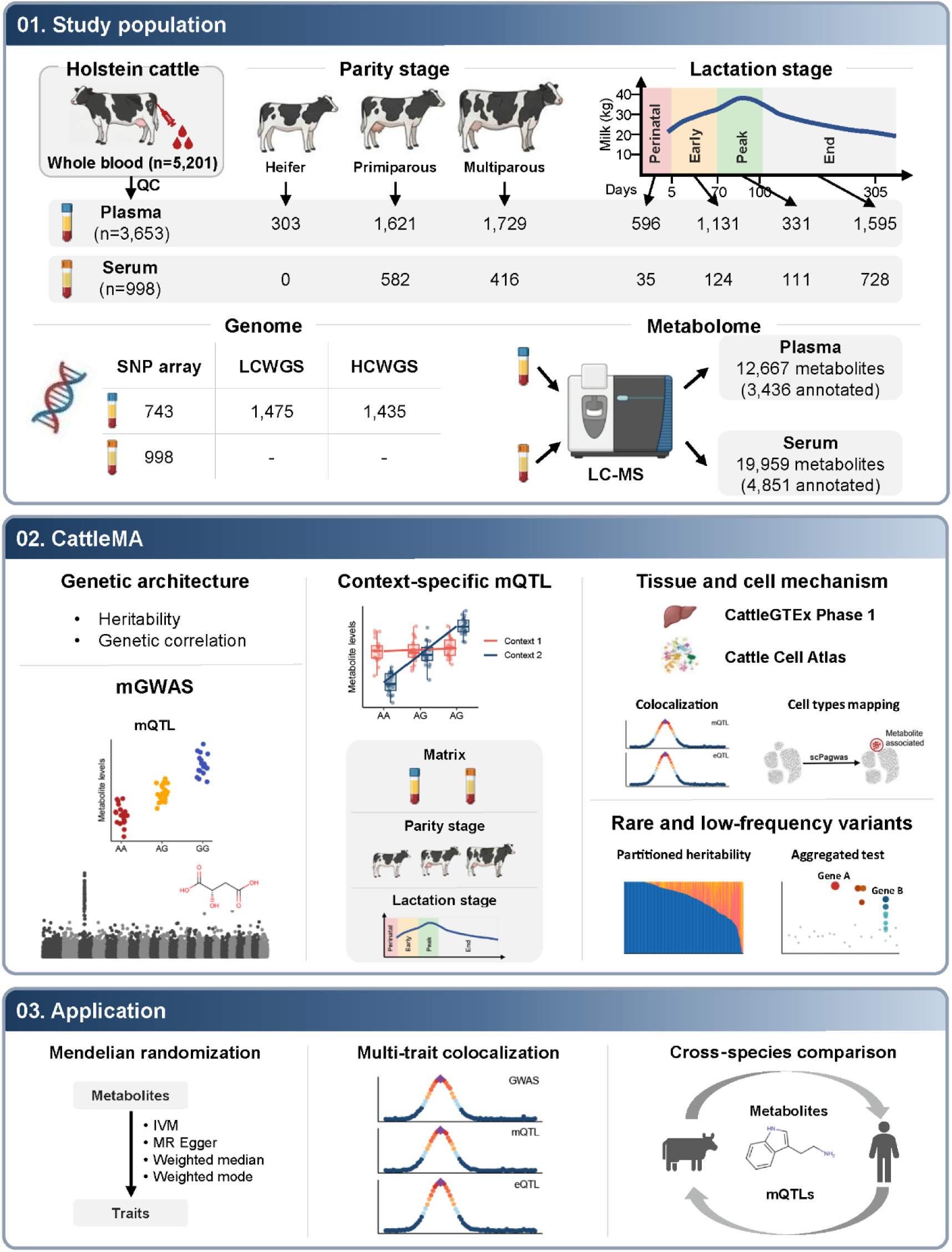
Overview of the CattleMA study.

## Results

### Data summary

We collected 5,206 blood samples from 4,990 Holstein cows (**Supplementary Table** 1). After quality control (**Supplementary Fig.** 1a, Methods), 4,651 high-quality samples were retained, comprising 3,653 plasma samples and 998 serum samples. These samples covered multiple physiological stages, including parity stages (heifer, primiparous and multiparous) and lactation stages (perinatal, early, peak and late lactation). All samples were genotyped using one of three platforms (**Supplementary Fig.** 1b). 150K SNP-array genotypes from 1,741 samples and LCWGS genotypes with average sequencing depth of 2.5× from 1,475 samples were imputed to sequence level using the multi-breed reference panel described in the Methods, whereas 1,435 samples were directly genotyped by HCWGS with average sequencing depth of 12.6×. For common variants (minor allele frequency (MAF) > 0.05), LCWGS-based imputation consistently outperformed SNP-array-based imputation, with an average accuracy (Pearson’s *r*) of 0.97 compared with 0.94 for SNP-array-based imputation (**Supplementary Fig.** 1c). After filtering out variants with MAF < 0.05 or Hardy-Weinberg equilibrium (HWE) test *P*-value < 1×10^-6^, a total of 9,061,760 SNPs was retained for downstream analyses.

Metabolite profiling of all samples was conducted using LC-MS, detecting 12,667 metabolic features in plasma and 19,959 in serum. Of these, 3,436 plasma metabolites (27%) and 4,851 serum metabolites (24%) were functionally annotated and included in subsequent analyses (**Supplementary Fig.** 1d and 1e; **Supplementary Table** 2). Among the annotated metabolites, 408 in plasma and 518 in serum were further assigned to 13 metabolic super-pathways, including amino acid, lipid, and nucleotide metabolism. A total of 2,482 metabolites overlapped between plasma and serum (**Supplementary Fig.** 1f). Across all metabolites, clustering analysis revealed that samples were separated primarily by plasma and serum (**Supplementary Fig.** 1j), rather than by parity or lactation stage (**Supplementary Fig.** 1k and 1l).

### Heritability and genetic correlation of metabolites

We first estimated the SNP-based heritability for metabolites measured in plasma and serum (**Supplementary Table** 3 and 4). Plasma heritability estimates ranged from 1 × 10^-6^ to 0.47, and 20.3% of metabolites were significantly heritable after Bonferroni correction (*P*-value < 0.05/3,436 = 1.5 × 10^-5^; median heritability = 0.0949; **Fig.** 2a). Median heritability varied across metabolic pathways, with higher estimates for lipid and cofactors/vitamins, and lower estimates for nucleotide. Serum metabolites showed higher heritability overall, ranging from 1 × 10⁻⁶ to 0.68; 18.2% were significant with a median heritability of 0.2286 (*P*-value < 0.05/4,851 = 1.0 × 10^-5^; **Fig.** 2b). Among 226 overlapping metabolites significant in both plasma and serum, heritability estimates showed a significant positive correlation (Pearson’s *r* = 0.67, *P*-value = 2.47 × 10⁻^22^; **Fig.** 2c).

**Fig 2.**
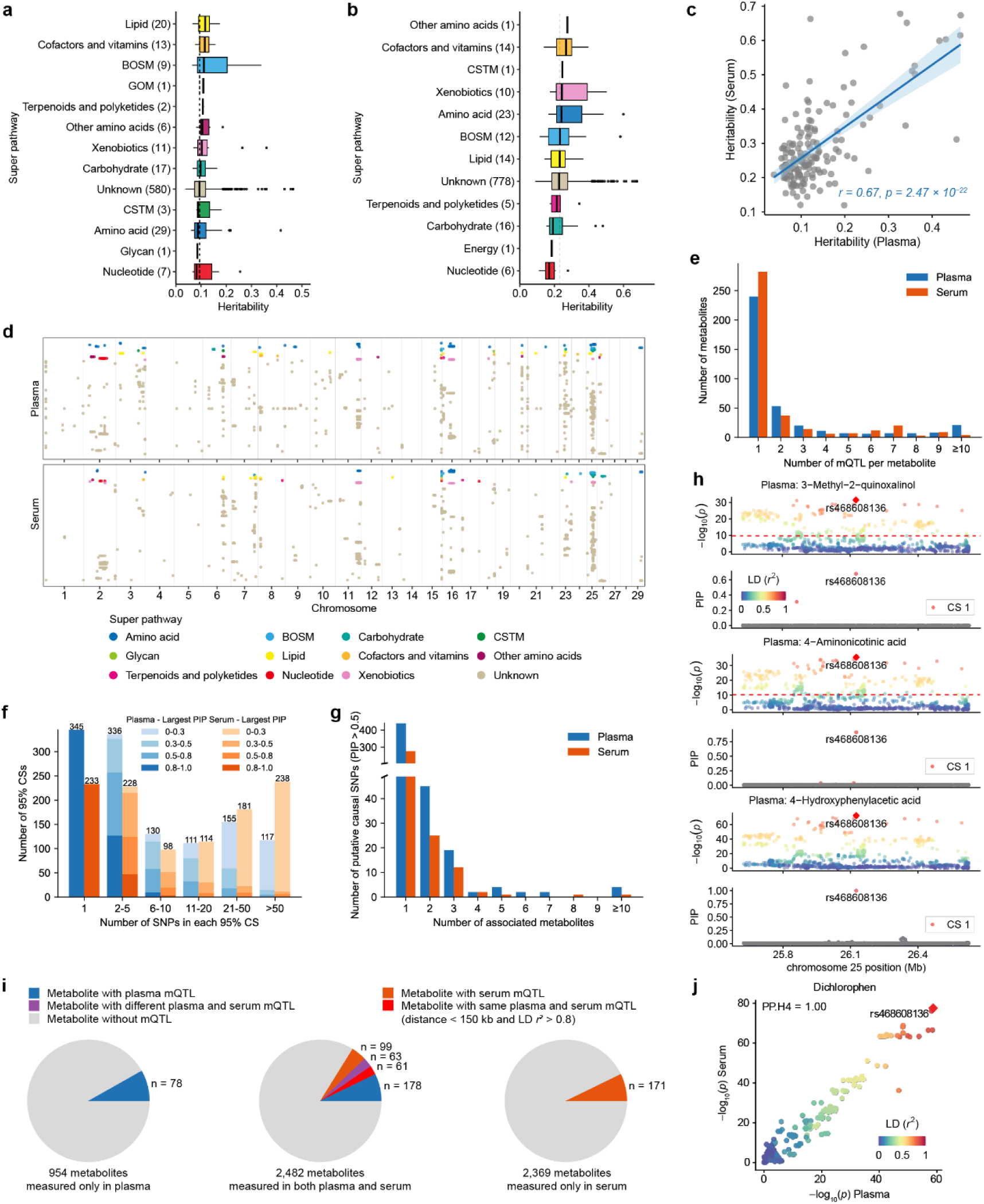
Genetic architecture of metabolites and mQTL discovery. **a**, **b**, Significant heritability estimates for plasma (**a**) and serum (**b**) metabolites across different super pathways. BOSM: Biosynthesis of other secondary metabolites; CSTM: Chemical structure transformation maps; GOM: Global and overview maps. Comparison of heritability of overlapping metabolites. **c**, Comparison of heritability estimates for metabolites measured in both plasma and serum. Pearson’s correlation coefficient and the corresponding *P*-value are shown. **d**, Summary Manhattan plots of mQTL associations in plasma (top) and serum (bottom). Each row represents a metabolite and is colored according to its super pathway. **e**, Distribution of the number of mQTL identified per metabolite. **f**, Number of variants in the 95% credible sets (CSs) and distribution of the posterior inclusion probabilities (PIPs) of the variants with the highest PIP within each CS. **g**, Distribution of the number of metabolites associated with putative causal variants (PIP > 0.5). **h**, Fine-mapping of mQTL for three plasma metabolites on chromosome 25. Regional association and fine-mapping plots are shown for 3-Methyl-2-quinoxalinol (top), 4-Aminonicotinic acid (middle), and 4- Hydroxyphenylacetic acid (bottom). For each metabolite, the upper panel displays Manhattan plot of GWAS results. Points colored in salmon represent the variants included in the CS of SuSiE. The lower panels show the PIP for each SNP as determined by SuSiE. The lead variant rs468608136 (indicated by text label) consistently shows the highest PIP across all three metabolites. **i**, Proportions and counts of metabolites with mQTL split by measurement matrix. **j**, Colocalization of the dichlorophen mGWAS signals between plasma and serum. Variants are colored according to their linkage disequilibrium (LD, *r*²) with the lead SNP.

To investigate the shared genetic architecture among metabolites, we estimated pairwise genetic correlations for all metabolite pairs within plasma or serum (**Supplementary Fig.** 2 and 3). In plasma, only 5.8% of metabolite pairs showed significant genetic correlations (*P*-value < 0.05/5,901,330 = 8.5 × 10^-9^), whereas this proportion was higher in serum, reaching 12.8% (*P*-value < 0.05/11,763,675 = 4.3 × 10^-9^). For example, strong positive genetic correlations were observed between D-(+)- glucose and D-mannose 6-phosphate (*r*_g_ = 0.99, *P*-value < 1 × 10^-300^), both involved in the carbohydrate metabolism, and between cholic acid and chenodeoxycholic acid (*r*_g_ = 0.94, *P*-value < 1 × 10^-300^) involved in the lipid metabolism. By contrast, D-(+)- glucose and L(-)-fucoseboth classified within the carbohydrate metabolism, showed a strong negative genetic correlation (*r*_g_ = -0.64, *P*-value = 7.7 × 10^-53^), suggesting potentially opposing genetic regulation among metabolites within the same broad metabolic pathway.

**Fig 3.**
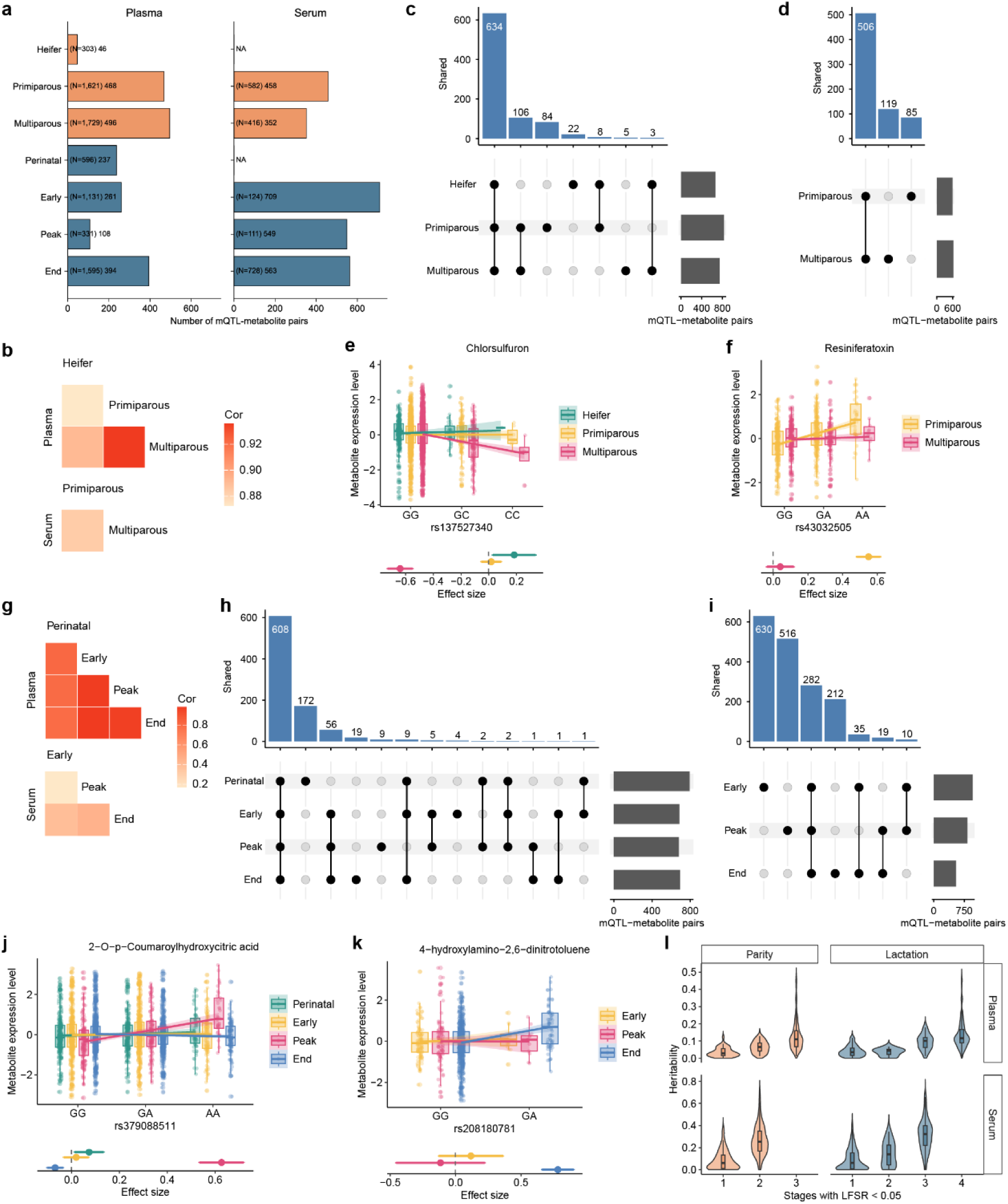
Context-dependent genetic regulation of metabolites. **a**, Numbers of mQTL-metabolite pairs identified at each stage, with the corresponding stage-specific sample sizes (N) shown in parentheses. **b**, Heatmap showing the Spearman’s correlation of mQTL effect sizes between parity stages, where effect sizes were estimated using MashR. **c** and **d**, UpSet plots showing the number of mQTL-metabolite pairs shared across the three parity stages in plasma (**c**) and serum (**d**). **e** and **f**, Examples of parity-stage-specific mQTLs for cobimetinib in plasma (**e**) and resiniferatoxin in serum (**f**). **g**, Heatmap showing the Spearman’s correlation of mQTL effect sizes between lactation stages, where effect sizes were estimated using MashR. **h** and **i**, UpSet plots showing the number of mQTL-metabolite pairs shared across the four lactation stages in plasma (**h**) and serum (**i**). **j** and **k**, Examples of lactation-stage-specific mQTLs for 2-O-p-coumaroylhydroxycitric acid in plasma (**j**) and 4-Hydroxylidocaine in serum (**k**). **l**, Heritability estimates for metabolites with mQTL shared across different numbers of parity or lactation stages, with shared associations defined using a MashR LFSR < 0.05.

### GWAS of plasma and serum metabolites

We performed mGWAS for plasma and serum metabolites using a linear mixed model implemented in GMAT. Genomic inflation factor analysis showed no evidence of systematic test statistic inflation due to population stratification (median λ = 0.997 for plasma and 0.989 for serum; **Supplementary Fig.** 4a and 4b). Among 3,436 and 4,851 plasma and serum metabolites being tested, we identified 414,115 and 470,933 significant associations for 380 and 395 metabolites (**Supplementary Fig.** 5a; **Supplementary Table** 5), respectively, with Bonferroni correction (*P* < 1.1 × 10^-10^ for plasma and *P* < 2.2 × 10^-10^ for serum, **Supplementary Fig.** 4c and 4d).

**Fig 4.**
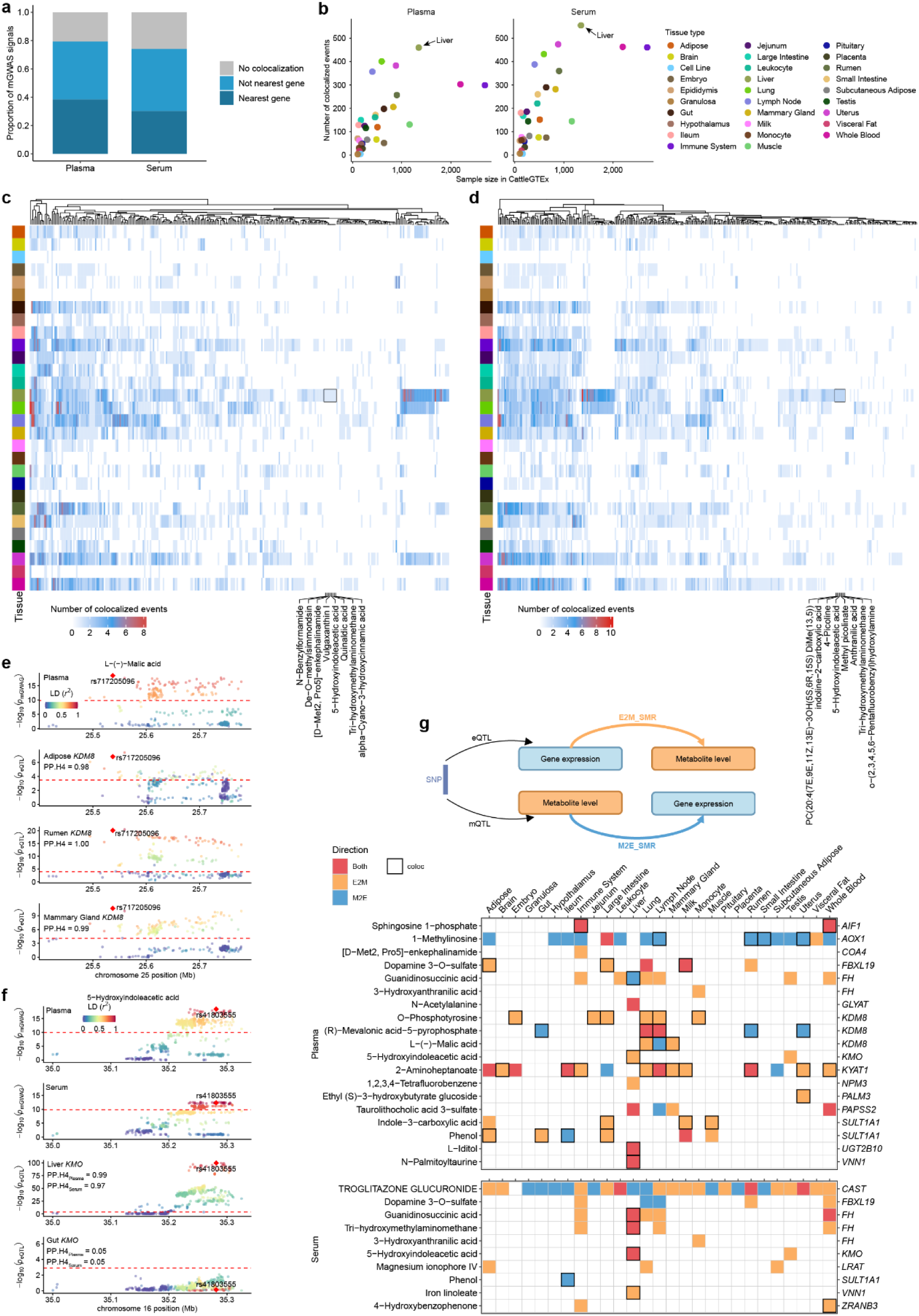
Associations of metabolite level with gene expression across tissues. **a**, Proportion of three types of mGWAS signals regarding the colocalization results. No colocalization, mGWAS loci that are not colocalized with any eGenes in 29 tissues. Not nearest gene, mGWAS loci whose colocalized eGenes are not nearest genes to mQTL. Nearest gene, mGWAS loci whose colocalized eGenes are the nearest ones. **b**, Relationship between number of colocalized events and sample size across 29 tissues in CattleGTEx. **c** and **d**, Heatmaps showing the numbers of colocalization events for plasma (**c**) and serum (**d**) metabolites across tissues. Rows represent tissues and columns represent metabolites. Metabolites are ordered by hierarchical clustering based on the tissue distribution of colocalization events. The colored bar on the left denotes tissue identity, corresponding to the tissues shown in **b**. Metabolites colocalized exclusively in liver are labeled. **e**, Regional colocalization plots for the plasma mQTL of L-(-)-malic acid and *cis*-eQTL for *KDM8* across three tissues: adipose, rumen and mammary gland. **f**, Regional colocalization plots for the plasma and serum mQTL of 5-hydroxyindoleacetic acid and *cis*-eQTL for *KMO* across two tissues: liver and gut. Variants are colored according to linkage disequilibrium (*r*²) with the lead colocalized variant. Dashed horizontal lines indicate the significance threshold. **g**, Direction of putative causal relationships between gene expression and metabolite level inferred from SMR analysis across tissues. Top, schematic illustration of the two possible directions tested by SMR: gene expression influencing metabolite level (E2M_SMR) and metabolite level influencing gene expression (M2E_SMR). Bottom, summary matrix of significant SMR associations between plasma or serum metabolites and *cis*-eQTL across tissues (*P*-valueSMR < 5 × 10^-6^ and *P*-valueHEIDI > 0.05). Rows represent metabolites and columns represent tissues; the corresponding candidate genes are shown on the right. Colored squares indicate the inferred direction of association: red, both directions supported; orange, gene expression to metabolite (E2M); blue, metabolite to gene expression (M2E). Black boxes denote metabolite-gene pairs supported by colocalization analysis (PP.H4 > 0.8).

For each metabolite with significant associations, we further detected conditionally independent SNPs using GCTA-COJO^34^. Overall, we identified 728 mQTL for 380 plasma metabolites and 622 mQTL for 395 serum metabolites (**Fig.** 2d; **Supplementary Table** 6). Among metabolites with mQTL, most of them harbored one mQTL (63% for plasma and 72% for serum; **Fig.** 2e), although a small number of metabolites had more than ten mQTL (21 for plasma and 4 for serum). Across the four metabolite annotation levels (Levels 1-4, as described in the Methods), the proportion of metabolites with at least one mQTL was broadly comparable, ranging from 9% to 17% in plasma and from 7% to 13% in serum (**Supplementary Fig.** 5b). The mQTL-associated metabolites were distributed across most super pathways, including 11 of 13 in plasma and 10 of 13 in serum (**Supplementary Fig.** 5c). Notably, 7 metabolites assigned to two super-pathways, the energy and global and the overview maps, showed no mQTL in either plasma or serum. Additionally, metabolites with mQTL exhibited significantly higher heritability than those without mQTL (*P*-value = 2.3 × 10^-52^ for plasma and *P*-value = 1.5 × 10^-86^ for serum; **Supplementary Fig.** 5d). The number of mQTL per metabolite was positively correlated with metabolite heritability (Pearson’s *r* = 0.54 for plasma and Pearson’s *r* = 0.50 for serum; **Supplementary Fig.** 5e).

To identify putative causal variants underlying mQTL, we created ±500 kb regions centered on each mQTL and merged overlapping regions associated with the same metabolite for fine-mapping analysis, resulting in 807 regions for plasma and 692 regions for serum. Within these regions, we detected putative causal variants in the majority of regions (96.9% for plasma and 96.5% for serum) (**Supplementary Fig.** 7a; **Supplementary Table** 7 and 8). For plasma metabolites, we detected 1,194 credible sets (CSs, PIP > 95%) of putative causal variants, including 1-402 variants (median = 4), of which 345 CSs consisted of a single variant (**Fig.** 2f). For serum metabolites, we detected 1,092 CSs of putative causal variants, ranging from 1 to 905 variants (median = 10), including 233 single-variant CSs (**Fig.** 2f). We considered individual variants with PIP > 0.5 as high-confidence candidate causal variants, including 522 variants in plasma across 715 CSs and 318 variants in serum across 403 CSs. Most metabolites were associated with a single putative causal variant (62% in plasma and 54% in serum; **Supplementary Fig.** 7b), and the majority of these variants were located in intergenic or intronic regions (68% in plasma and 71 % in serum; **Supplementary Fig.** 7c and 8d). In plasma, 45 putative causal variants were shared by at least two metabolites (up to 20; **Fig.** 2g). For example, rs468608136 showed PIP of 0.68, 0.91 and 1.00 for 3- methyl-2-quinoxalinol, 4-aminonicotinic acid, and 4-hydroxyphenylacetic acid, respectively (**Fig.** 2h). Similarly, in serum, 42 putative causal variants were shared by at least two metabolites (**Fig.** 2g).

We next compared mQTL discovery across metabolites measured in plasma, serum or in both matrices. An mQTL was detected for 8% of plasma-specific metabolites (78 of 954), 7% of serum-specific metabolites (171 of 2,369), and 16% of metabolites measured in both plasma and serum (401 of 2,482; **Fig.** 2i). Among these 401 cross-matrix metabolites with an mQTL detected in at least one matrix, 277 (69%) showed a significant association in only one matrix, including 178 detected exclusively in plasma and 99 detected exclusively in serum. The remaining 124 metabolites had mQTL detected in both plasma and serum. Of these, 61 metabolites (49%) were associated with the same or highly correlated mQTL across matrices, defined by a distance of < 150 kb and LD *r*² > 0.8 (**Fig.** 2i). To determine whether these signals reflected shared causal variants, we further performed pairwise colocalization analysis and identified 49 metabolites with strong evidence of shared causal variants between plasma and serum (PP.H4 > 0.8; **Supplementary Table** 9). For example, an mQTL (rs468608136) showed conserved regulatory effects on dichlorophen levels in both plasma and serum (PP.H4 = 1; **Fig.** 2j). The direction and effect size for 56 colocalized mQTL were nearly identical in both plasma and serum (**Supplementary Fig.** 7). Additionally, we applied MashR^35^ all 1,048 mQTL of 401 overlapping metabolites, which accounted for the statistical power of mQTL discovery due to sample size differences. At the local false sign rate (LFSR) threshold of 0.05, 1,168 mQTL-metabolite pairs were significant. Among them, approximately 63.8% were shared between plasma and serum, whereas 24.2% appeared plasma-specific and 12.0% serum-specific (**Supplementary Fig.** 8a). For example, rs207824937 was strongly associated with N(6)-(1-carboxyethyl)-L-lysine levels in plasma (*P*-value_mGWAS_ = 2.6 × 10^-20^, MAF = 0.15), but showed no evidence of association in serum (*P*-value_mGWAS_ = 0.72, MAF = 0.14) (**Supplementary Fig.** 8b). Similarly, rs109672550 was associated with lysosulfatide levels specifically in serum (**Supplementary Fig.** 8c).

### The dynamic landscape of genetic effects on metabolites across parity and lactation stages

To investigate context-dependent mQTL regulation, we first performed context-stratified mGWAS analysis within three parity stages and four lactation stages, followed by conditional analysis using GCTA-COJO^34^ to identify independent mQTL. Across all 12 context-stratified analyses, only 1-6% of metabolites were associated with at least one = (**Fig.** 3a), and the proportion of metabolites with detectable mQTL increased when sample size increased (**Supplementary Fig.** 9a and 9b). We then applied MashR to all mQTL-metabolite pairs, including 995 parity-stratified pairs in plasma and 801 in serum, as well as 979 lactation-stratified pairs in plasma and 1,817 in serum, to characterize context-specific mQTL effects.

For parity-dependent mQTL regulation, effect sizes were highly correlated across parity stages, particularly between primiparous and multiparous cows in plasma (Pearson’s *r* = 0.94; **Fig.** 3b) and between the two evaluated parity stages in serum (Pearson’s *r* = 0.88; **Fig.** 3b). Accordingly, most mQTL-metabolite pairs were shared across parity stages, with 74% of plasma signals shared across multiple stages and 71% of serum signals shared between both stages (**Fig. 3c** and 3d). Nevertheless, 13% of plasma signals and 29% of serum signals were detected in only one parity stage. For example, rs43381709 showed a significant association with plasma cobimetinib specifically in heifers (**Fig.** 3e), whereas rs43032505 was associated with serum esiniferatoxin specifically in primiparous stage (**Fig.** 3f).

For lactation-dependent mQTL regulation, mQTL effects were generally concordant among lactation stages, particularly among early, peak, and end lactation, where plasma effect-size correlations exceeded 0.97 (**Fig.** 3g). Most plasma mQTL-metabolite pairs were shared across multiple lactation stages (68%), whereas 23% were restricted to a single stage (**Fig.** 3h). Context-specific associations were also observed in serum, a substantial number of stage-restricted associations (80%; **Fig.** 3i). Representative examples included rs379088511, which was associated with plasma 2-O-P- coumaryldihydroxycitric acid specifically during peak lactation (**Fig.** 3j), and rs382917416, which showed a peak-lactation-specific association with serum 4- hydroxylidocaine (**Fig.** 3k).

Across both plasma and serum, metabolites whose mQTL were shared across a greater number of parity or lactation stages generally exhibited higher heritability (**Fig.** 3l). Overall, we identified a subset of mQTL signals with clear context-specific regulatory effects on plasma and serum metabolites.

### Exploring the multi-tissue molecular architecture underlying circulating metabolites

To explore the potential regulatory mechanisms of plasma and serum metabolites, we integrated mGWAS signals with *cis*-eQTL from 29 tissues in CattleGTEx Phase 1^32^ Colocalization analysis revealed that 290 of 380 (79%) tested plasma metabolites and 344 of 395 (74%) tested serum metabolites were significantly colocalized with at least one *cis*-eQTL (PP.H4 > 0.8; **Fig.** 4a). Notably, among these colocalizations, 41% of plasma and 44% of serum mGWAS signals were not colocalized with the nearest genes to mQTL, indicating the complexity of gene regulation underlying metabolites. Across 29 tissues, liver showed the largest number of colocalization events in both plasma and serum (**Fig.** 4b), consistent with its key role in regulating circulating metabolite levels^36,37^.

In total, we identified 4,469 colocalization events corresponding to 2,098 gene-metabolite pairs in plasma (**Supplementary Table** 10), and 5,791 colocalization events corresponding to 2,275 gene-metabolite pairs in serum (**Supplementary Table** 11). The number of colocalization events per metabolite was positively correlated with the number of mQTL per metabolite in both plasma and serum (Pearson’s *r*_plasma_ = 0.38 and Pearson’s *r*_serum_ = 0.37, respectively; **Supplementary Fig.** 10a). Most metabolites showed colocalization with *cis*-eQTL across multiple tissues (**Fig. 4c and 4d**), including 272 of 290 plasma metabolites and 311 of 344 serum metabolites, with a median of 8 tissues in both plasma and serum (**Supplementary Fig.** 10b and 10c). These results indicate that plasma and serum metabolite levels are subject to widespread regulation across tissues. Nevertheless, a subset of metabolites showed tissue-specific colocalization, with signals restricted to a single tissue for 18 plasma metabolites and 33 serum metabolites. For example, 8 plasma metabolites and 8 serum metabolites colocalized exclusively with liver (**Fig. 4c** and 4d). As expected, the implicated genes by colocalization were significantly enriched in multiple metabolism-related Gene Ontology (GO) terms and KEGG pathways, such as organic acid metabolic process, oxoacid metabolic process, and retinol metabolism pathways (**Supplementary Fig.** 10d and 10e).

We provided several colocalization examples below: The plasma mQTL of L-(-)-malic acid showed colocalization with eQTL of *KDM8* at rs717205096 across 17 tissues, including adipose, rumen, mammary gland, and other tissues (**Fig.** 4e). This broad pattern is biologically plausible, as malic acid is a core intermediate of the tricarboxylic acid cycle (**Supplementary Fig.** 11), known as a central metabolic pathway active across multiple tissues. *KDM8*, a member of the histone demethylase family, has been reported to regulate *PKM2*^38^, a key glycolytic enzyme; glycolysis feeds into the tricarboxylic acid cycle through the conversion of pyruvate to acetyl-CoA, thereby linking glucose catabolism to mitochondrial energy metabolism. The mQTL of 5- Hydroxyindoleacetic acid were colocalized with *cis*-eQTL of *KMO* at rs41803555 only in the liver, in both plasma (PP.H4 = 0.99) and serum (PP.H4 = 0.97), illustrating a tissue-specific colocalization pattern (**Fig.** 4f). KMO is a key enzyme in the kynurenine pathway, the predominant route of tryptophan catabolism, which accounts for ∼90% of tryptophan metabolism and takes place largely in the liver (**Supplementary Fig.** 12) ^39^. 5-hydroxyindoleacetic acid is a second pathway of tryptophan metabolism and *KMO* may reflect coordinated regulatory mechanisms within the tryptophan metabolic network. Consistently, three additional metabolites from serum in tryptophan metabolism, 3-hydroxyanthranilic acid, anthranilic acid, and kynurenic acid, showed strong colocalization (PP.H4 > 0.96) with *cis*-eQTL of *KMO* at rs41803555 in the liver (**Supplementary Fig.** 13).

Furthermore, we performed stage-stratified colocalization analysis to prioritize shared putative causal variants between stage-mQTL and *cis*-eQTL. The tissue distribution of regulatory signals was broadly consistent across parity and lactation stages in both plasma and serum, with the liver contributing a substantial fraction of colocalization events across stages (mean = 9.1%; **Supplementary Fig.** 14a). Most metabolites colocalized in multiple tissues, consistent with the pattern observed in the merged analysis (**Supplementary Fig.** 14b), indicating that the tissue-specific regulatory effects on metabolite levels are largely preserved across different stages. Of note, 26 plasma metabolites and 53 serum metabolites colocalized only within one stage with one gene in one tissue, highlighting clear context-specific regulatory effects (**Supplementary Fig.** 14c and 15). For example, at the *KMO* locus in the liver, the colocalization signal for 5-hydroxyindoleacetic acid was detected specifically in the lactation stage, illustrating stage-dependent genetic regulation of this metabolite in the liver (**Supplementary Fig.** 14d).

### Putative causal relationships between gene expression and metabolite levels

Colocalization analysis indicates a shared putative causal variant likely regulating two traits, but the effect direction cannot be established by itself. Variants regulating gene expression (eQTL) may influence metabolite levels, whereas variants affecting metabolite levels (mQTL) may, in turn, influence gene expression. We therefore performed summary data-based Mendelian randomization (SMR) analysis using *cis*- eQTL and mQTL summary statistics, considering gene expression as the exposure and metabolite levels as the outcome (represented as E2M_SMR), or metabolite levels as the exposure and gene expression as the outcome (represented as M2E_SMR), followed by the heterogeneity in dependent instruments (HEIDI) test to differentiate a causal or pleiotropic model from a linkage model^40^ (**Fig.** 4g).

In plasma, we identified 2,427 E2M_SMR associations corresponding to 398 genes and 239 metabolites, and 1,561 M2E_SMR associations corresponding to 261 genes and 226 metabolites (*P*-value_SMR_ < 5 × 10^-6^ and *P*-value_HEIDI_ > 0.05), with 502 shared associations in both E2M_SMR and M2E_SMR (**Supplementary Fig.** 16a; **Supplementary Table** 12 and 13). In serum, we identified 3,062 E2M_SMR associations corresponding to 398 genes and 275 metabolites, and 1,996 M2E_SMR associations corresponding to 217 genes and 255 metabolites (*P*-value_SMR_ < 5 × 10^-6^ and *P*-value_HEIDI_ > 0.05), with 609 shared associations in both E2M_SMR and M2E_SMR (**Supplementary Fig.** 16a; **Supplementary Table** 14 and 15). Cross-matrix comparison showed that 479 E2M associations were shared between plasma and serum, representing 19.8% and 15.7% of plasma and serum E2M signals, respectively; 243 M2E associations were shared, representing 15.6% and 12.2% of plasma and serum M2E signals, respectively. A larger number of significant E2M_SMR associations compared to M2E_SMR suggests that the genetic effect of gene expression on metabolite levels is a more predominant mechanism than the genetic effect of metabolite levels on gene expression. Consistent with the colocalization results, most metabolites showed significant SMR associations across multiple tissues in both plasma and serum (**Supplementary Fig.** 16d-g).

In total, 30-36% of SMR signals were supported by colocalization, including 30% of plasma E2M_SMR, 34% of plasma M2E_SMR, 30% of serum E2M_SMR, and 36% of serum M2E_SMR associations (**Supplementary Fig.** 16c). For instance, colocalization analysis detected a shared genetic basis between *KDM8* expression in the mammary gland and plasma L-(-)-malic acid levels, and the significant plasma E2M_SMR signal (*P*-value_SMR_ = 8.85 × 10^-8^ and *P*-value_HEIDI_ = 0.10) further suggested that *KDM8* expression may influence plasma L-(-)-malic acid levels (**Fig.** 4g). Similarly, plasma and serum 5-hydroxyindoleacetic acid colocalized with liver *KMO*, with significant E2M_SMR signals detected in both plasma (*P*-value_SMR_ = 1.29 × 10^-15^ and *P*-value_HEIDI_ = 0.12) and serum (*P*-value_SMR_ = 4.91 × 10^-12^ and *P*-value_HEIDI_ = 0.26), whereas an additional M2E_SMR signal was observed in serum (*P*-value_SMR_ = 5.07 × 10^-12^ and *P*-value_HEIDI_ = 0.25), suggesting a more complex and potentially bidirectional relationship.

### Detecting metabolite-relevant cell types

To detect the cell types relevant with metabolites, we integrated cattle single-cell transcriptomic data from the CattleCA^33^, comprising 130 cell types across seven major cell lineages in 59 tissues (**Supplementary Table** 16). To prioritize cell types potentially involved in metabolic regulation, we performed a cell type-metabolism enrichment analysis using scPagwas^41^, revealing distinct associations between specific cell lineages and metabolic pathways (**Fig.** 5a). For example, stromal cells were predominantly associated with carbohydrate metabolism and the biosynthesis of secondary metabolites, whereas nervous cells showed significant enrichment for several amino acid metabolic pathways (**Fig.** 5a, c). Overall, stromal cells exhibited significantly stronger enrichment across nearly all metabolic categories, highlighting their central roles in metabolic regulation, consistent with previous reports^42^ (**Fig.** 5b).

**Fig 5.**
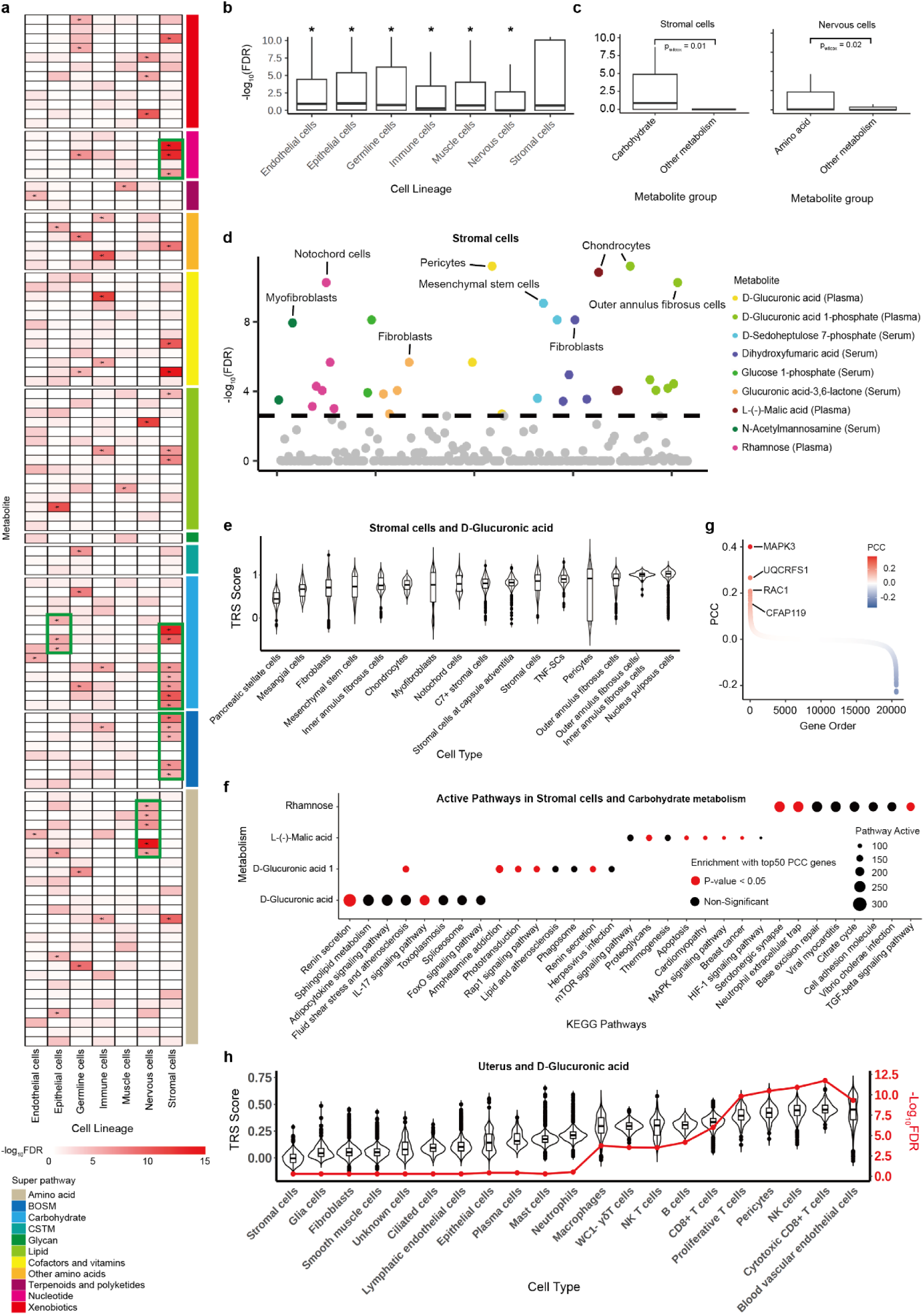
Discovery of metabolite-relevant cell types. **a**, Association between cell lineages and metabolisms. The heatmap represents the significant level of each cell lineage with metabolisms. The asterisk represents significant association. **b.** Global association between each cell lineage and all metabolisms. **c**, Association between stromal cells and nervous cells with carbohydrate metabolism and amino acid metabolism. **d**, Association between carbohydrate metabolism and stromal cells. Each circle represents a cell type–trait association. The *x* axis represents cell lineages sorted alphabetically. **e**, Association between stromal cells and D-Glucuronic acid. The violin plot presents the TRS for each cell type with the D-Glucuronic acid. **f**, Active KEGG pathways in the top significant trait-relevant cell types. Dot size represents the significance level (-log_10_(*P*)) of pathway activity; the dot color represents the significance level (FDR) after multiple comparisons of the enrichment between active pathway genes and top metabolism-relevant genes in the given cell type. **g**, PCC distribution of genes in pericytes with D-Glucuronic acid. **h**, Association between cell types in uterus and D-Glucuronic acid. The violin plot presents the TRS for each cell type with the D-Glucuronic acid. The red line represents the significance level (-log_10_(*P*)) for each cell type with the D-Glucuronic acid under a two-sided hypergeometric test.

To further investigate stromal cell-specific metabolic regulation, we examined associations between 16 stromal cell types and nine carbohydrate metabolic pathways. Among these, pericytes and chondrocytes displayed the highest enrichment and trait-relevant score (TRS) compared with other stromal cell types. Specifically, pericytes were preferentially linked to D-glucuronic acid metabolism, whereas chondrocytes were enriched for both L-(-)-malic acid and D-glucuronic acid 1-phosphate metabolism (**Fig.** 5d, e). Notably, the top 50 genes with high Pearson’s correlation coefficients (top 50 PCCs) for D-glucuronic acid levels in pericytes were significantly enriched in the IL-17 signaling pathway (**Fig.** 5f). *MAPK3*, the gene with the highest PCC value (**Fig.** 5g), is a key regulator of multiple metabolic processes^43^, including substrate phosphorylation during glucose metabolism^44^, suggesting that pericytes may participate in D-glucuronic acid regulation through inflammatory and metabolic signaling networks associated with IL-17 activity. Consistently, pericytes also exhibited elevated enrichment significance and TRS scores in uterus tissue, where they were specifically enriched (**Fig.** 5h). Moreover, the colocalized eGene *CFAP119* in the uterus showed a positive correlation with D-glucuronic acid levels (PCC = 0.15; **Fig.** 5g; **Supplementary Table** 17), indicating that pericytes may mediate the observed genetic association between *CFAP119* and the mQTL involving D-glucuronic acid in the uterus. Similarly, chondrocytes exhibited the highest TRS scores for D-glucuronic acid 1- phosphate metabolism, with *MAPK3* again representing the top correlated gene with the highest PCC value (**Supplementary Fig.** 17a, b), suggesting a shared regulatory mechanism underlying carbohydrate metabolism across different stromal cell types. In addition, the top 50 PCC-associated genes for L-(-)-malic acid in chondrocytes were significantly enriched in the proteoglycan-related pathways and the MAPK signaling pathway, both previously implicated in glucose metabolic regulation^45,46^ (**Fig.** 5f).

Furthermore, we performed enrichment analysis between 13 nervous cell types and 12 amino acid metabolic pathways. Among these cell types, glial cells showed the strongest enrichment significance for 3-hydroxyanthranilic acid metabolism **(Supplementary Fig.** 17c). Genes with high PCC values (top 50 PCC genes) for 3- hydroxyanthranilic acid levels in glial cells were significantly enriched in the glutamatergic and dopaminergic synapse pathways, suggesting potential regulation of glial cells on 3-hydroxyanthranilic acid metabolism through neurotransmitter-associated metabolic and synaptic signaling processes^47^. Notably, *AKT3*, the gene with the highest PCC value, is a key regulator of neuronal signaling and synaptic plasticity and has been implicated in glutamatergic and dopaminergic neurotransmission^48^, further supporting the potential role for glial cells in the neurotransmitter-associated metabolic regulation of 3-hydroxyanthranilic acid (**Supplementary Table** 18).

### Rare and low-frequency variants reveal additional genetic architecture underlying metabolites

Given the limited imputation accuracy of rare and low-frequency variants (MAF < 0.05) from SNP array data (*r* = 0.80) and the substantially higher accuracy achieved with LCWGS (*r* = 0.97; **Supplementary Fig.** 1c), we analyzed LCWGS and HCWGS data from 2,910 individuals with plasma metabolite profiles, retaining 16,834,909 high-quality SNPs after QC to quantify the contribution of non-common variants to metabolite variation. Variants were stratified into rare (MAF < 0.005), low-frequency (0.005 ≤ MAF < 0.05) and common (MAF ≥ 0.05) classes, which accounted for 26.2%, 22.6%, and 51.2% of all variants, respectively (**Fig.** 6a). Among 575 metabolites with significant SNP heritability estimated using all variants (*P*-value < 0.05/3,436 = 1.5 × 10^-5^), incorporating rare and low-frequency variants explained slightly but significantly more heritability than analyses restricted to common variants alone (median = 0.1047 vs 0.1037; *P*-value = 2.6 × 10^-20^; **Fig.** 6b), indicating a measurable contribution from rare and low-frequency variants to metabolite variation. Although common variants accounted for the majority of heritability for these metabolites (mean = 77%), rare and low-frequency variants contributed substantially to a subset of metabolites, with rare and low-frequency variants individually explaining more than 50% of the heritability for 14 and 45 metabolites, respectively (**Fig.** 6c).

**Fig. 6.**
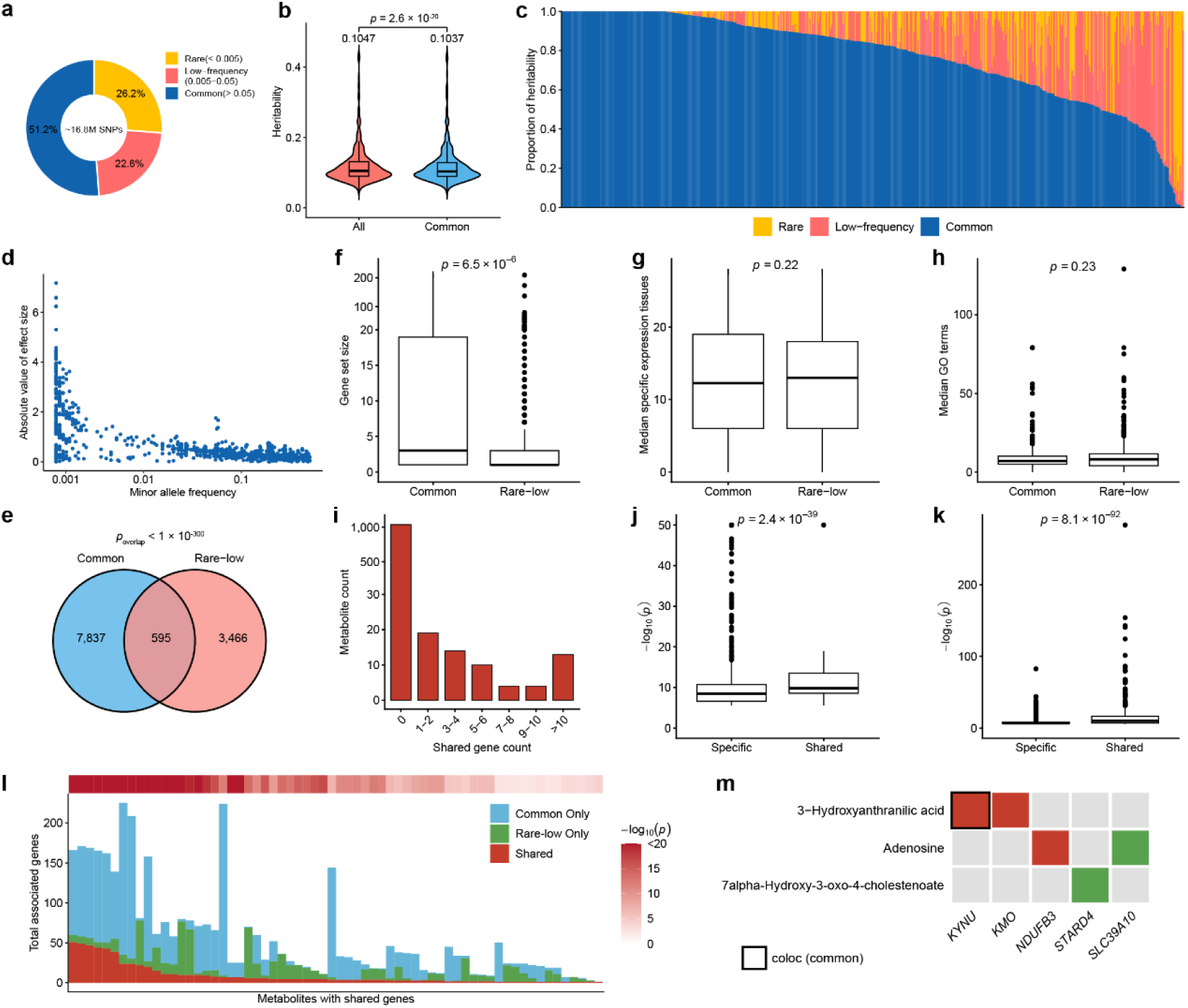
Contribution and convergence of rare, low-frequency and common variants to genetic architecture of metabolite. **a**, Proportion of variants stratified by minor allele frequency (MAF) into rare (MAF < 0.005), low-frequency (0.005 ≤ MAF < 0.05), and common (MAF ≥ 0.05). **b**, Distribution of metabolite heritability with significant heritability estimates (*P*-value < 0.05/3436 = 1.5 × 10^-5^) calculated using all variants or common variants only. Numbers above the violin plots indicate median heritability. *P*-value was calculated using a paired Wilcoxon signed-rank test. **c**, Proportion of metabolite heritability explained by rare, low-frequency, and common variants across metabolites with significant heritability estimates (*P*-value < 0.05/3436 = 1.5 × 10^-5^). Each vertical bar represents one metabolite, ordered by the proportion of heritability explained by common variants. **d**, Relationship between absolute effect size and MAF of mQTL. **e**, Overlap of gene-metabolite pairs with significant genes in common-variant and rare/low-frequency frequency variant gene-based analyses. *P*-value above the Venn diagrams indicates the significance of overlap, calculated using one-sided hypergeometric tests. **f**, Comparison of gene set size between common-variant and rare/low-frequency variant associated genes. Significance of overlapping was evaluated by hypergeometric test. **g**, Comparison of tissue specificity, measured as the median number of tissues with gene expression, between common-variant and rare/low-frequency variant associated genes. **h**, Comparison of functional annotation breadth, measured as the median number of Gene Ontology (GO) terms, between common-variant and rare/low-frequency variant associated genes. **i**, Distribution of the number of shared genes per metabolite between common-variant and rare/low-frequency variant analyses. **j**, **k**, Comparison of association strength between specific and shared genes in the common-variant analyses (**j**) and the rare/low-frequency variant analyses (**k**). **l**, Number of associated genes per metabolite among metabolites with overlapping genes between common-variant and rare/low-frequency variant analyses, stratified into common-only, rare/low-frequency-only and shared genes. The top heatmap shows the significance of shared-gene enrichment based on a hypergeometric test. **m**, Examples of genes implicated by common-variant and rare/low-frequency variant analyses. The color scale is the same as that used in panel **l**.

We next performed mGWAS using all variants. Consistent with previous human study^9^, low-frequency mQTL exhibited larger effect sizes than common mQTL (**Fig.** 6d). Given the limited power of single-variant mGWAS for rare and low-frequency variants, we further conducted gene-based association using the aggregated Cauchy association test for variant-level *P*-values (ACAT-V) method^49^. For comparison, common-variant gene-based associations were tested using MAGMA^50^. In total, 19,462 protein-coding genes were tested for common variants and 19,970 protein-coding genes were tested for rare and low-frequency variants. At Bonferroni-corrected significance thresholds (*P*-value < 0.05/19,462 = 2.6 × 10^-6^ for common variants and *P*-value < 0.05/19,970 = 2.5 × 10^-6^ for rare and low-frequency variants), we identified 4,062 significant common-variant gene-metabolite associations (**Supplementary Table** 19) and 8,432 significant rare/low-frequency variant gene-metabolite associations (**Supplementary Table** 20), of which 595 were shared between the two testes (*P*-value < 1 × 10^-300^; **Fig.** 6e). Rare and low-frequency variant-associated genes formed significantly smaller gene sets per metabolite than common-variant associated genes (*P*-value = 6.5 × 10^-6^; **Fig.** 6f), whereas tissue specificity and functional annotation breadth were comparable between them (*P*-value = 0.22 and *P*-value = 0.23, respectively; **Fig.** 6g and 6h).

We further assessed convergence between common and rare/low-frequency signals at the gene level. 46 metabolites shared at least one associated gene between common-variant and rare/low-frequency variant analyses, including 16 metabolites sharing more than ten genes (**Fig.** 6i). Shared gene-metabolite pairs exhibited significantly stronger association evidence than specific gene-metabolite pairs in both common-variant and rare/low-frequency variant analyses (*P*-value = 2.4 × 10^-39^ and *P*-value = 8.1 × 10^-92^, respectively; **Fig.** 6j and 6k). Moreover, 58 metabolites exhibited significantly greater shared genes than expected by chance (FDR-adjusted *P*-value < 0.05; **Fig.** 6l), although shared genes represented only a subset of the associated genes for these metabolites (median proportion: 28% for common-variant associations and 50% for rare/low-frequency variant associations). Representative convergent examples included *KYNU*/*KMO* and 3-hydroxyanthranilic acid, consistent with the direct roles of *KYNU* and *KMO* in the kynurenine pathway of tryptophan degradation (**Supplementary Fig.** 12); the *KYNU* signal was further supported by *cis*-eQTL colocalization (**Fig.** 6m). We also identified rare/low-frequency-specific associations, such as *STARD4* and 7α- hydroxy-3-oxo-4-cholestenoate. Given the implicated role of *STARD4* in intracellular cholesterol trafficking^51^ and the function of 7α-hydroxy-3-oxo-4-cholestenoate as an intermediate in the conversion of cholesterol to bile acids, this association highlights the additional biological insights provided by rare/low-frequency variants. Together, these findings demonstrate that rare and low-frequency variants uncover complementary metabolite-associated genes beyond those identified by common variants alone, thereby expanding the allelic spectrum of genetic architecture underlying the cattle metabolites.

### Understanding the metabolic and molecular mechanisms underlying complex traits

To investigate the putative causal effects of metabolites on complex traits, we performed two-sample MR analysis by integrating our metabolite data with GWAS summary statistics from 11 Holstein cattle complex traits, including five milk production traits, three reproduction traits, and three health traits^52–54^ (**Supplementary Table** 21). By applying four MR methods (inverse variance weighted (IVW), MR-Egger, weighted median, and weighted mode) in the TwoSampleMR package^55^, we identified 198 putative causal metabolite-trait associations (FDR-adjusted *P*-value <

0.05 in at least two of the four MR methods, with *P*-value > 0.05 for both heterogeneity and horizontal pleiotropy tests) in plasma, involving 99 metabolites and 10 traits, and 213 putative causal metabolite-trait associations in serum, involving 126 metabolites and 11 traits (**Supplementary Fig.** 17a; **Supplementary Table** 22 and 23). Among trait-associated metabolites, 55 of 99 in plasma (56%) and 58 of 126 in serum (46%) showed putative casual effects on multiple traits (**Supplementary Fig.** 17b). Notably, 45 associations were shared between plasma and serum (*P*-value = 1.05 × 10^-9^; **Supplementary Fig.** 17c). For example, plasma dehydroepiandrosterone sulfate showed a putative causal effect on protein percentage (PP; *β[SE]*= 0.015200.0144;; **Supplementary Fig.** 17d). This metabolite is a precursor of steroid hormones^56^ (e.g., estrogens), which have key roles in mammary gland development and lactation^57^. Similarly, serum thyroxine sulfate showed a putative causal effect on heifer conception rate (HCR; *β[SE]*= 0.091700.0144;; **Supplementary Fig.** 17e). Thyroxine sulfate is a major thyroid hormone metabolite that plays a key role in fertility and embryo development^58^.

To further investigate the molecular mechanisms underlying complex traits, we integrated *cis*-eQTL and mQTL with 532 GWAS signals of these 11 cattle traits. Overall, GWAS signals showed stronger enrichment in mQTL regions than in whole-blood *cis*- eQTL regions, with the strongest enrichment observed for plasma mQTL regions (**Fig.** 7a). Further comparison revealed that more than 20% of GWAS loci overlapped with mQTL across multiple trait categories (**Fig.** 7b). We next performed multi-trait colocalization analysis to determine shared causal variants among trait GWAS loci, mQTL, and *cis*-eQTL. Among the 148 GWAS loci overlapping plasma mQTL, 23 (16%) showed evidence of colocalization with both mQTL and *cis*-eQTL (PPA.abc > 0.5), whereas 8 were supported only by mQTL colocalization (**Fig.** 7c). Among the 140 GWAS loci overlapping serum mQTL, 14 (10%) were colocalized with both mQTL and *cis*-eQTL, whereas one was supported only by mQTL colocalization (**Fig.** 7c). Notably, GWAS variants shared by *cis*-eQTL and mQTL explained a greater proportion of phenotypic variance than mQTL-specific in both plasma and serum (**Fig.** 7d). In total, we identified 539 significant gene-metabolite-trait trios in plasma (**Supplementary Table** 24), involving 99 genes and 32 metabolites, and 102 significant gene-metabolite-trait trios in serum (**Supplementary Table** 25), involving 38 genes and 24 metabolites. Liver and mammary gland contributed a relatively large number of gene-metabolite pairs for milk production traits, particularly protein percentage (**Fig.** 7e and 7f). Several gene-metabolite pairs were colocalized with multiple traits; for example, brain-*LYNX1* and plasma kaempferol 3-O-rhamnoside showed colocalization across six traits (**Fig.** 7g). Most gene-metabolite-trait pairs were specific, but 14 were detected in both plasma and serum (*P*-value = 7.97 × 10^-28^; **Fig.** 7h), and their corresponding multi-trait colocalization signals showed a moderate positive correlation in PPA.abc values, the posterior probability of colocalization among GWAS, mQTL and *cis*-eQTL (Pearson’s *r* = 0.47, *P*-value = 9.31 × 10^−2^; **Fig.** 7i).

**Fig 7.**
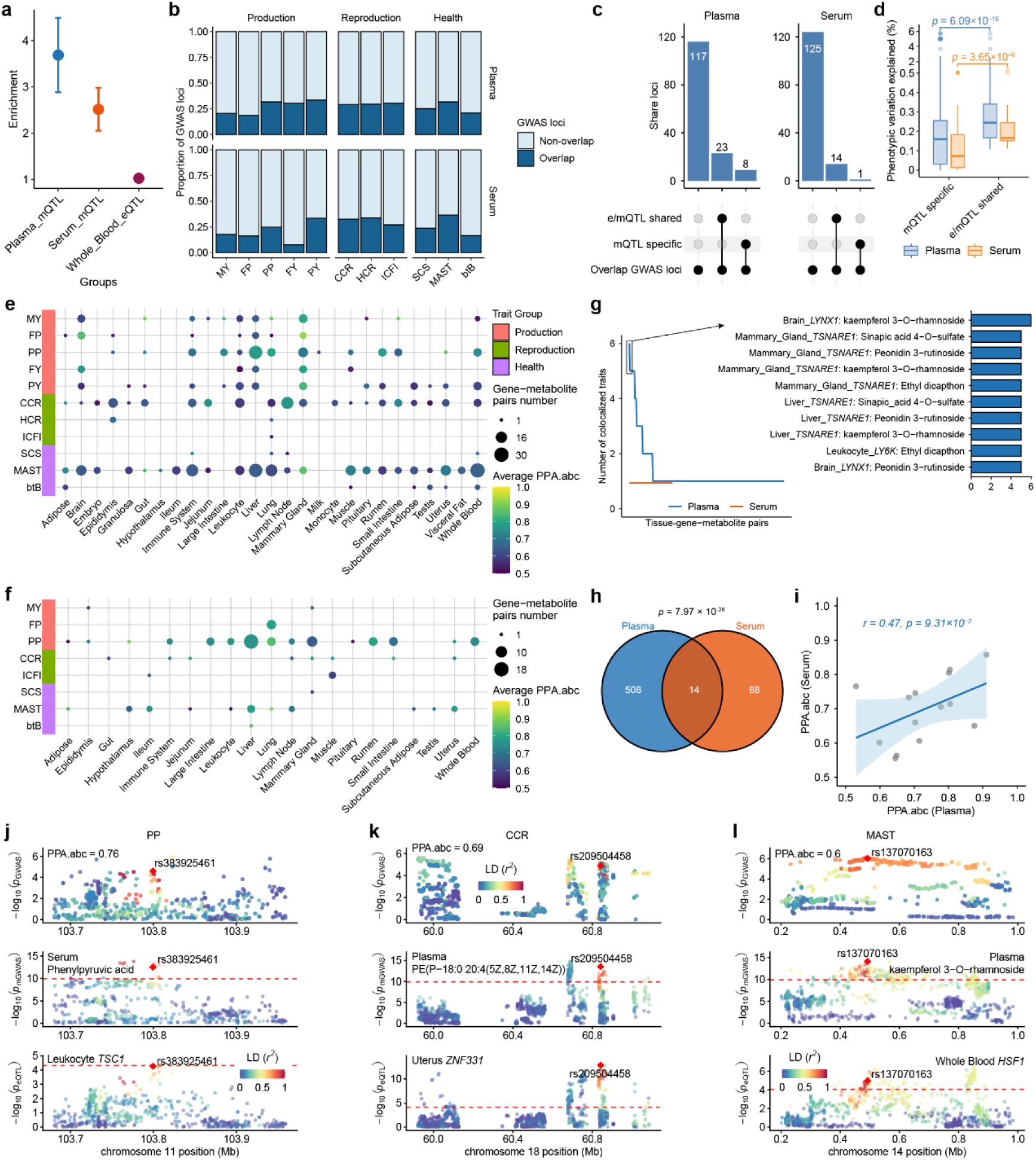
Integrative analysis of *cis*-eQTL, mQTL, and trait GWAS signals. **a**, Enrichment analysis of GWAS signals in plasma mQTL regions, serum mQTL regions, and whole-blood *cis*-eQTL regions. **b**, Proportion of GWAS loci for production, reproduction, and health traits that overlap with plasma or serum mQTL regions. **c**, UpSetR plot depicting the overlap between GWAS hits across colocalization types, i.e., e/mQTL-shared: GWAS signals are colocalized with mQTL and *cis*-eQTL. mQTL-specific: GWAS signals are only colocalized with mQTL. **d**, Phenotypic variation explained by mQTL-specific and e/mQTL-shared loci. Statistical significance was determined using two-sided Wilcoxon rank-sum test. **e**, Number of gene-metabolite (plasma) pairs that were colocalized (PPA.abc > 0.5) with GWAS loci of complex traits across tissues. **f**, Number of gene-metabolite (serum) pairs that were colocalized (PPA.abc > 0.5) with GWAS loci of complex traits across tissues. Point size indicates the number of gene-metabolite pairs, whereas point color indicates the average PPA.abc of each colocalization. **g**, Number of colocalized traits per tissue-gene-metabolite pair in plasma and serum. The right panel highlights representative gene-metabolite pairs with colocalization more than five traits. **h**, Overlap of tissue-gene-metabolite-trait pairs identified in plasma and serum. *P*-value above the Venn diagrams indicate the significance of overlap, calculated using one-sided hypergeometric tests. **i**, Correlation of PPA.abc values for tissue-gene-metabolite-trait pairs shared between plasma and serum. The solid line indicates the linear regression fit, and the shaded area denotes the 95% confidence interval. Pearson’s correlation coefficient and two-sided *P*-value are shown. **j**, Regional colocalization plots for PP GWAS, serum phenylpyruvic acid mQTL, and leukocyte *TSC1 cis*-eQTL. **k**, Regional colocalization plots for CCR GWAS, plasma PE(P-18:0/20:4(5Z,8Z,11Z,14Z)) mQTL, and uterus *ZNF331 cis*-eQTL. **l**, Regional colocalization plots for MAST GWAS, plasma kaempferol 3-O-rhamnoside mQTL, and whole-blood *HSF1 cis*-eQTL. Variants are colored according to their linkage disequilibrium (*r*²) with the lead colocalized variant. Dashed horizontal lines indicate the significance threshold.

Three representative examples further illustrated these integrative relationships. For milk production traits, the GWAS signal of protein percentage colocalized with the serum phenylpyruvic acid mQTL and leukocyte *TSC1 cis*-eQTL at rs383925461 (PPA.abc = 0.76; **Fig.** 7j). Phenylpyruvic acid is an intermediate of phenylalanine metabolism that regulates milk protein synthesis via the LAT1-mTOR signaling pathways in bovine mammary epithelial cells^59^, whereas *TSC1* is an inhibitor of mTOR signaling and is significantly upregulated during lactation^60^. Although a direct regulatory link between *TSC1* and phenylalanine remains unclear, there may be a potential interaction between metabolic and transcriptional regulation underlying PP. For reproduction traits, the GWAS signal of cow conception rate (CCR) colocalized with the plasma PE (P-18:0/20:4(5Z,8Z,11Z,14Z)) mQTL and uterus *ZNF331 cis*-eQTL at rs209504458 (PPA.abc = 0.69; **Fig.** 7k). This metabolite belongs to a class of glycerophospholipids, which play a pivotal role in embryonic development^61^. *ZNF331*, a zinc-finger transcriptional regulator, is predicted to regulate *PISD*, supported by three databases (GTRD^62^, hTFtarget^63^, and ChIPBase^64^; **Supplementary Fig.** 19), which encodes a key enzyme in PE (P-18:0/20:4(5Z,8Z,11Z,14Z)) biosynthesis. Together, these results suggest a potential regulatory link in which uterus *ZNF331* may influence CCR through effects on PE (P-18:0/20:4(5Z,8Z,11Z,14Z)). For health traits, the GWAS signal of mastitis colocalized with the plasma kaempferol 3-O-rhamnoside mQTL and whole-blood *HSF1 cis*-eQTL at rs137070163 (PPA.abc = 0.60; **Fig.** 7l). Kaempferol 3- O-rhamnoside has been reported to exhibit anti-inflammatory and anti-oxidant properties^65^, whereas *HSF1* is a stress-responsive transcription factor linked to breast cancer in humans^66^ and heat tolerance, milk fat, and milk protein in cattle^67,68^. Together, these results demonstrate that integrating e/mQTL-GWAS colocalization signals provides deeper insight into the molecular mechanisms underlying complex traits.

### Comparative regulatory effects on metabolites between cattle and humans

To assess the cross-species conservation of metabolite genetic regulation, we compared metabolite heritability estimates and the contribution of orthologous regulatory loci between cattle and humans. We collected publicly available human mGWAS summary statistics for metabolites that matched those profiled in our study, based on HMDB ID or metabolite name (**Supplementary Table** 26 and 27). In total, we obtained 37 overlapping plasma metabolites with mGWAS summary statistics from the Canadian Longitudinal Study on Aging (CLSA; *n* = 8,299)^69^, and 38 overlapping serum metabolites from a meta-analysis of 2,901 Han Chinese individuals across three cohorts-the Zhejiang Metabolic Syndrome Cohort (ZMSC, *n* = 1,124), the Westlake Precision Birth Cohort (WeBirth, *n* = 908), and the Tongji-Huaxi-Shuangliu Birth Cohort (THSBC, *n* = 869)^70^. We investigated the concordance of metabolite heritability estimates between cattle and humans. Overall, metabolite heritability showed limited concordance between cattle and humans. A significantly positive correlation was observed only for serum metabolites in the WeBirth cohort (Pearson’s *r* = 0.69, *P*-value = 1.2 × 10^-3^), whereas no significant correlation was observed for plasma metabolites in the CLSA cohort (Pearson’s *r* = 0.25, *P*-value = 0.13) or for serum metabolites in the ZMSC cohort (Pearson’s *r* = 0.13, *P*-value = 0.51) and THSBC cohort (Pearson’s *r* = 0.18, *P*-value = 0.42) (**Supplementary Fig.** 20a and 20b). This limited and heterogeneous concordance may partly reflect differences in cohort composition, sample size, environmental exposures, metabolomics platforms^71,72^.

To further determine whether human mQTL aid in understanding the genetic regulation of cattle metabolites due to evolutionary constraints, we compared the proportion of heritability explained by loci orthologous to significant human mGWAS loci (*P*-value < 1 × 10^-5^) with that explained by MAF-matched random loci. Using UCSC LiftOver, 43% of plasma mQTL and 46% of serum mQTL were successfully mapped to orthologous cattle loci (**Fig.** 8a). For both plasma and serum metabolites, orthologous loci corresponding to significant human mGWAS signals generally explained a greater proportion of heritability than MAF-matched random loci (*P*-value = 0.01 for plasma and *P*-value = 0.04 for serum; **Fig. 8b and 8c**), supporting partial cross-species conservation of regulatory effects on metabolites. Among the 37 overlapping plasma metabolites, orthologous human mGWAS loci explained more than 5% of their heritability for 14 metabolites and more than 50% of heritablity for four metabolites, with indolelactic acid showing the highest proportion value (74%; **Supplementary Fig.** 20c). Similarly, among the 38 overlapping serum metabolites, orthologous loci explained more than 5% of heritability for 11 metabolites and more than 50% of heritability for three metabolites, with stearamide showing the highest proportion value (99.9%; **Supplementary Fig.** 20d). For example, a pair of orthologous variants- rs2736163 in humans and rs133318407 in cattle-showed conserved regulatory effects on sphingosine 1-phosphate in both species (*P*-value_Human_ = 2.45 × 10^-6^ and *P*-value_Cattle_ = 2.70 × 10^-8^; **Fig.** 8d). Both variants reside within *PRRC2A* in cattle and humans, which encodes a recently identified N6-methyladenosine (m⁶A) reader that directly targets *CSNK1E* to enhance WNT signaling^73^, a pathway with known functional interactions with sphingosine-1-phosphate signaling^74^.

**Fig 8.**
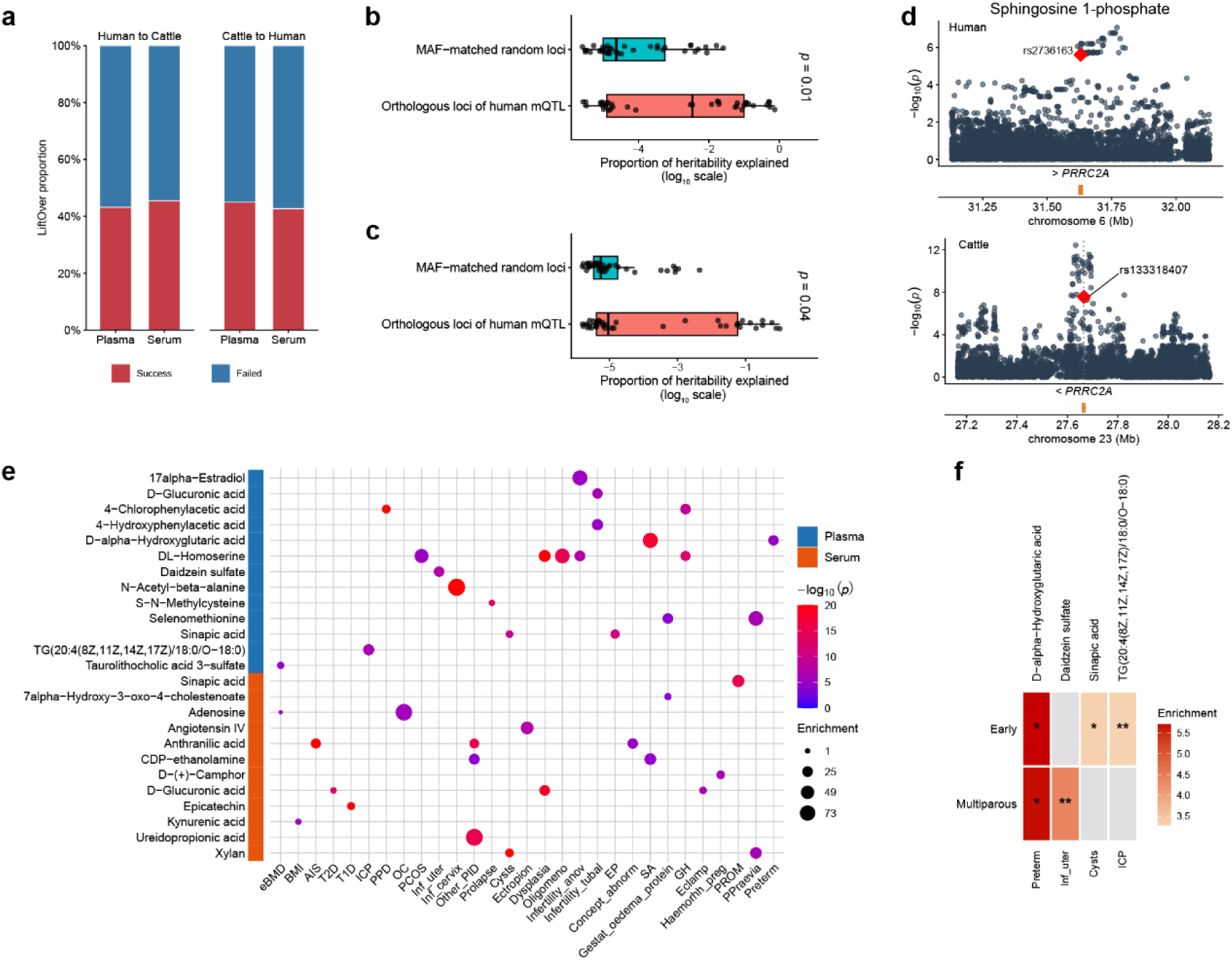
Cross-species comparison of orthologous mQTL between cattle and humans. **a,** Proportion of human mQTL successfully mapped to the cattle genome, and cattle mQTL successfully mapped to the human genome. **b**, **c**, Proportion of SNP heritability explained by different loci groups for overlapping plasma (**b**) and serum (**c**) metabolites. Statistical significance was assessed using two-sided Wilcoxon rank-sum test. **d**, Comparison of *P*-values for orthologous loci regulating sphingosine 1-phosphate between cattle and human mGWAS. Example of a conserved plasma mQTL for sphingosine 1-phosphate in cattle and humans, where human variant rs2736163 is orthologous to cattle variant rs133318407 (ARS-UCD1.2). **e**, Partitioned heritability enrichment of orthologous loci of cattle plasma and mQTL across representative human traits and diseases. **f**, Heatmap showing significant partitioned heritability enrichment of stage-dependent orthologous cattle mQTL in human female reproductive health traits. Rows indicate cattle physiological stages, and columns indicate metabolite-trait combinations. Color intensity represents enrichment, and asterisks indicate statistical significance levels (*: *P*-value < 0.05; **: *P*-value < 0.01).

Finally, we evaluated whether cattle mQTL contribute to human complex trait variation. We performed stratified linkage disequilibrium score regression (S-LDSC)^75^ using orthologous cattle mQTL annotations and GWAS summary statistics for 56 human traits and diseases, spanning aging, metabolism, immune response categories, and female reproductive health (**Supplementary Table** 28). Overall, 45% of fine-mapped plasma mQTL (7,340 loci for 307 metabolites) and 43% of fine-mapped serum mQTL (10,669 loci for 363 metabolites) were successfully mapped to orthologous human loci (**Fig.** 8a). After Bonferroni correction for each trait (*P*-value < 0.05/307 = 1.6 × 10^-4^ for plasma metabolites and *P*-value < 0.05/363 = 1.4 × 10^-4^ for serum metabolites), we identified 205 significant plasma metabolite-trait pairs (**Supplementary Table** 29), involving 106 plasma metabolites and 52 human traits, together with 240 significant serum metabolite-trait pairs (**Supplementary Table** 30), involving 132 serum metabolites and 53 human traits. These enrichments included known or biologically plausible metabolite-trait relationships (**Fig.** 8e), such as adenosine with estimated bone mineral density (eBMD), TG(20:4(8Z,11Z,14Z,17Z)/18:0/O-18:0) with intrahepatic cholestasis of pregnancy (ICP), sinapic acid with ovarian cyst (Cysts), daidzein sulfate with inflammatory disease of uterus (Inf_uter), and D-alpha-Hydroxyglutaric acid with preterm labor and delivery (Preterm). These examples are consistent with previous studies implicating adenosine in eBMD^76^, lipid metabolic dysregulation in ICP^77^, sinapic acid in ovarian metabolic dysfunction and PCOS-related ovarian fibrosis^78^, and daidzein-derived metabolites in estrogen receptor-mediated and inflammatory processes in endometrial disease^79,80^. Although direct evidence linking D-alpha- hydroxyglutaric acid to preterm birth remains limited, metabolomic studies have broadly implicated metabolic perturbations in spontaneous preterm labor and delivery^81^.

Furthermore, we tested whether regulatory effects shaped by cattle physiological stage specificity could advance our understanding of human complex traits and diseases. Among the four pregnancy- and postpartum-related traits identified above, stage-stratified S-LDSC analyses revealed that orthologous cattle mQTL active at specific physiological stages were preferentially enriched for several female reproductive health traits (**Fig.** 8f). In particular, early-lactation-dependent orthologous mQTL were significantly enriched with ICP through TG(20:4(8Z,11Z,14Z,17Z)/18:0/O-18:0) (enrichment = 3.29, *P*-value = 1.1 × 10^-3^). Likewise, multiparous-stage-specific orthologous mQTL were significantly enriched for Inf_uter through daidzein sulfate (enrichment = 5.67, *P*-value = 0.03), which is supported by previous evidence that multiparity has been suggested as a risk factor for endometritis^82^. Together, these findings demonstrate the evolutionary conservation of blood metabolite regulatory architecture and highlight the cattle metabolome as a valuable comparative framework for interpreting the genetic basis of human complex traits and diseases.

## Discussion

The genetic architecture of circulating metabolites provides insights into the biological complexity of metabolic regulation and its contribution to complex traits. Here, we established the CattleMA, a comprehensive resource comprising 4,651 circulating blood samples, including 3,653 plasma and 998 serum samples with 3,436 and 4,851 confidently annotated metabolites, respectively. The CattleMA resource identified 728 independent mQTL for 380 plasma metabolites and 619 independent mQTL for 395 serum metabolites. By integrating whole-genome sequence variants with plasma and serum metabolomics, bulk and single-cell datasets, and complex-trait GWAS, CattleMA moves beyond mQTL discovery towards a mechanistic view of metabolic regulation. A central finding is that the genetic architecture of circulating metabolites comprises both stable and dynamic components. More than 60% of mQTL effects were shared across parity and lactation stages, indicating a broadly conserved genetic basis of metabolism across physiological states. However, the presence of context-specific mQTL demonstrates that a subset of genetic effects is modulated by the marked metabolic and endocrine transitions accompanying reproductive maturation and lactation^83^. Together, these findings indicate that otherwise latent genetic effects can become apparent under specific physiological conditions.

Extensive colocalization between mQTL and *cis*-eQTL implicated transcriptional regulation as an important mechanism linking genetic variants to circulating metabolite abundance^4^, with more than 70% of mQTL signals showing evidence of colocalization with the liver emerging as a major tissue of action^2^. Single-cell integration further resolved specialized regulatory programs in stromal cell populations, particularly pericytes and chondrocytes, emphasizing that circulating metabolite levels reflect both tissue-level regulation and more restricted cell-type-specific processes^33,84^. By connecting genetic variants, gene expression, metabolites and complex traits, our analyses additionally support circulating metabolites as intermediate molecular phenotypes that can help bridge regulatory variation and complex traits. For example, the colocalization of the CCR GWAS signal with plasma PE (P- 18:0/20:4(5Z,8Z,11Z,14Z)) and the uterus *ZNF331 cis*-eQTL implicates coordinated regulation of glycerophospholipid metabolism in female fertility^61^. Although some associations do not establish a definitive causal cascade, they generate experimentally testable hypotheses and indicate that gene expression and circulating metabolites represent complementary layers through which genetic effects may influence complex traits.

Our study also investigated the contribution of rare and low-frequency variants to metabolic traits. Although common variants explained most metabolite heritability on average (∼77%), non-common variants made substantial contributions to certain metabolites, consistent with observations for human complex traits^85,86^. Aggregate testing identified 8,432 gene-metabolite associations, both reinforcing common-variant findings and revealing additional signals not captured by conventional analyses. These results provide a more comprehensive view of the allelic architecture underlying circulating metabolites in cattle.

Beyond mechanistic interpretation within cattle, our cross-species analyses revealed partial conservation of metabolite regulatory architecture between cattle and humans. Although overall metabolite heritability showed limited concordance across species, orthologous loci corresponding to significant human mGWAS loci explained more cattle metabolite heritability than matched random loci. Orthologous cattle mQTL regions were also enriched for the heritability of diverse human complex traits, including metabolic, inflammatory and reproductive disorders. Notably, some enrichments depended on cattle parity or lactation stage, indicating that physiologically activated regulatory programs in cattle may help prioritize disease-relevant pathways in humans. Together, these findings support livestock as a valuable comparative model for identifying evolutionarily conserved metabolic mechanisms while distinguishing context- and species-specific regulation^87–89^.

Despite these advances, several limitations still need further considerations and address for the development of CattleMA project in the future: 1) sample sizes across physiological stages were imbalance in stage-specific analysis, limiting statistical power for detecting stage-specific mQTL. 2) only common SNPs, low-frequency variants, and rare variants were used in mQTL analysis. Other types of genomic variants (e.g., structural variants and somatic mutations)^90–92^ can be incorporated to assess their effects on metabolite levels in mQTL analysis. Extended complementary molecular phenotypes, such as epigenomics, proteomics, long-read genomics and transcriptomics, and single-cell multi-omics, could be integrated to construct a more complete regulatory landscape underlyig metabolites; 3) only untargeted metabolomics technique were used for metabolite identification. Untargeted metabolomics enabled comprehensive metabolite profiling, but it provides predominantly relative quantification. Future efforts will integrate untargeted metabolomics with targeted metabolomics, improving absolute quantification, metabolite identification, and analytical reproducibility^93^. 4) novel metabolite features remain unannotated in untargeted metabolomics using small-molecule-associated peaks because of the limited coverage of existing spectral libraries. Recent advances in chemical language models will anticipate previously uncharacterized metabolites from untargeted mass spectrometry data, offering a powerful strategy to overcome the low annotation rate that remains a major limitation of large-scale metabolome atlas^94^. Using the expanded metabolome atlases, metabolomic language foundation models assist in disease risk estimation and chronic disease prediction, demonstrating their abilities in early disease prediction from routine blood-derived metabolic profiles^95,96^. 5) the metabolic states of individual cells remain largely unresolved. Emerging single-cell metabolomics technologies will enable to uncover cell-to-cell heterogeneous and spatial metabolism within tissues^24^. 6) Few biological contexts (parity stage and lactation stage) were considered in this study. Future releases of CattleMA will expand to additional biological contexts, including breed, sex, developmental stage, environmental exposure, nutrition factor, and disease status, following the guidelines proposed in the FarmGTEx White Paper^31^.

## Methods

### Ethics

All experimental procedures in this study were approved by the Animal Ethics Committee of the Shandong Academy of Agricultural Sciences, China. Ethical approval for animal use and survival experiments was granted by the Animal Ethics Committee (IASVM-2026-009).

### Sample collection

A total of 5,206 whole blood samples were collected from the tail vein of 4,990 Chinese Holstein cows from 14 dairy farms in Shandong Province, China. All blood sample collections occurred during the period that was after the first morning milking but before the first morning feeding. The sampled animals represented a range of physiological conditions spanning different parities and lactation stages, as detailed in **Supplementary Table** 1. Parity was classified as heifer (calving number = 0), primiparous (calving number = 1), and multiparous (calving number ≥ 2). Lactation stage was defined according to days in milk (DIM) as perinatal (DIM < 5), early (5 ≤ DIM < 70), peak (70 ≤ DIM < 100) and end lactation (DIM ≥ 100). The collected samples were processed in two cohorts, yielding 4,147 plasma samples and 1,059 serum samples.

### Data generation and processing

#### Metabolite data

For plasma sample collection from each individual, one 10-ml vacutainer blood collection tube containing the anticoagulant of heparin sodium was used. All collected plasma samples were centrifuged at 3,000 g for 10 minutes at 4 ℃, and 400 µL supernatant of them were immediately transferred to liquid nitrogen for 15-minute freezing. For serum sample collection from each individual, one 10-ml vacutainer blood collection tube containing the coagulant of silicon dioxide was used. All collected serum samples were placed at room temperature for 30 minutes and then centrifuged at 3000 g for 10 minutes at 4 ℃ as well. The 400 µL supernatant of them were also immediately transferred to liquid nitrogen for 15-minute freezing. The aforementioned plasma and serum samples were promptly sent to the Beijing Genomics Institute (BGI) on dry ice for metabolite identification using the untargeted metabolomics technique. All metabolite detection of plasma and serum samples were measured using the UPLC I-Class Plus (Waters, USA) tandem Q Exactive (QE) high-resolution mass spectrometer (Thermo Fisher Scientific, USA). A total of 12,667 metabolites were quantified in plasma samples and 19,959 metabolites in serum samples. Metabolite annotation was performed using three reference databases-mzCloud, BGI metabolome database (BMDB), and human metabolome database (HMDB)-using Compound Discoverer 3.3 software. Metabolite annotation results were classified into five levels (Level1-Leve5) as follows: 1. Level1 indicates metabolite annotation to the same precursor ion, secondary mass spectrum (i.e., fragment ion), and retention time to reference databases; 2. Level2 indicates metabolite annotation to the same precursor ion and secondary mass spectrum to reference databases, with the secondary mass spectrum score ≥ 60; 3. Level3 indicates metabolite annotation to the same precursor ion and secondary mass spectrum to reference databases, with the secondary mass spectrum score < 60; 4. Level4 indicates metabolite annotation to the same precursor ion only to reference databases; 5. Level5 indicates no metabolite annotation to any databases. After annotation, 3,436 plasma metabolites and 4,851 serum metabolites with assigned identities were retained for subsequent analyses.

#### Genotype data

Genomic DNA was extracted from whole blood of all individuals and used for genotyping. Three genotyping platforms were applied (**Supplementary Table** 1). DNA sequencing for LCWGS and HCWGS datasets was performed using the Salus Pro gene sequencer (Shenzhen Salus BioMed Co., Ltd.). A total of 1,872 individuals were genotyped using GGP Bovine HDv3 (150K). SNP coordinates were harmonized to the bovine reference genome (ARS-UCD1.2) using LiftOver (v1.3.3)^97^. Then, array-based genotypes were subsequently imputed to the whole-genome sequence level using Beagle (v5.1)^98^, with a reference panel comprising 3,530 cattle and 28,166,177 SNPs. Low-coverage whole-genome sequencing (LCWGS) was performed for 1,677 individuals at an average depth of ∼2.5×. Clean reads were mapped to the bovine reference genome (ARS-UCD1.2) using BWA-MEM (v0.7.17)^99^. File conversion, sorting, and indexing were conducted with SAMtools (v1.6)^100^, and PCR duplicates were removed using Picard tools (https://broadinstitute.github.io/picard/, v2.21.2). Then, we used GLIMPSE2 (v2.0.0)^101^ to impute genotype based on the same reference panel described above. High-coverage whole-genome sequencing (HCWGS) was conducted for 1,594 individuals at an average depth of ∼12.6×. Variant calling was performed using the Sentieon (v202503.01) DNASeq pipeline (https://www.sentieon.com/products/). Briefly, clean reads were aligned to the reference genome using the “bwa mem” algorithm, followed by BAM sorting (“util sort”) and duplicate removing (“Dedup”). Raw GVCFs were generated with “Haplotyper”, and joint genotyping across samples was carried out using “GVCFtyper” to produce cohort-level VCF files. SNPs were filtered using the “VariantFiltration” function in GATK (v4.0.7.0)^102^ with the following criteria parameters: “QD < 2.0||MQ < 40.0||FS > 60.0||SOR > 3.0||MQRankSum < -12.5||ReadPosRankSum < -8.0||QUAL < 30”. Then, we used Beagle to impute genotypes of these individuals with the same reference panel above. Finally, VCF files from all three data sources were merged using BCFtools (v1.21)^103^ to generate a unified genotype dataset for downstream analyses.

To evaluate imputation accuracy, we selected 100 cows from the HCWGS dataset with sequencing depth greater than 15× as validation individuals. Their sequence-level genotypes were used as the truth genotypes. To mimic SNP-array and LCWGS data, the HCWGS data were reduced either to the corresponding SNP-array marker set or by downsampling reads to 2.5×. The reduced datasets were then imputed back to sequence level using the same imputation pipeline as applied to the full dataset. To reduce computational time, we only performed imputation on chromosome 25. For each variant, imputation accuracy was quantified using two metrics: genotype concordance (GC), defined as the proportion of matching genotypes between imputed and true, and *r*, defined as the Pearson correlation between imputed expected dosages and true genotypes.

#### Data cleaning of quantified metabolites and genotypes

Samples were excluded if they were duplicates, had missing genotype data, had unknown parity, or had DIM > 500 or unknown. After filtering, 4,651 samples remained for downstream analyses, including 3,653 plasma samples and 998 serum samples. To ensure the accuracy and consistency of metabolomic measurements, quality control (QC) procedures were applied separately to the plasma and serum datasets to remove systematic bias and background noise. Following previous studies^69^, metabolite levels were natural log-transformed, extreme outliers (> 3 standard deviations from the mean) were excluded, and values were standardized to a mean of 0 and a standard deviation of 1. The imputed genotypes of these samples retained only variants with a minor allele frequency (MAF) ≥ 0.05 and *P*-value of Hardy-Weinberg Equilibrium test > 1×10^-6^, resulting in a final set of 9,061,760 SNPs for subsequent analyses.

### SNP-based heritability and genetic correlation estimation

A genomic relationship matrix (GRM) was constructed from all 9,061,760 common SNPs. SNP-based heritability for each plasma and serum metabolite was estimated using a linear mixed model implemented in GCTA software (v1.95.1)^104^. Genetic correlations between pairs of metabolites within plasma or serum were estimated using FastGC method implemented in OmiGA software (v2.0.3)^105^.

### Genome-wide association study of plasma and serum metabolites

We performed mGWAS for plasma and serum metabolites, separately, using GMAT software (https://github.com/chaoning/GMAT) with a linear mixed model, adjusting for herd, detection batch, parity, lactation stage, and the first two principal components (PCs). Genome-wide significance threshold was determined using a Bonferroni correction accounting for the effective number of both independent SNPs and independent metabolites. To calculate the effective number of independent SNPs, we performed LD pruning using PLINK (v1.90)^106^ with the command of “--indep-pairwise 50 5 0.2”, retaining variants with pairwise *r*^2^ < 0.2. After LD pruning, 365,576 SNPs remained. To account for correlations among metabolites, we calculated the effective number of independent metabolites by performing principal component analysis (PCA) on the metabolite matrices and retaining the number of PCs that cumulatively explained > 95% of the total variance^107,108^. In plasma, 1,322 PCs captured > 95% of the variance, whereas 628 PCs captured > 95% of the variance in serum (**Supplementary Fig.** 4b and 4c). Therefore, the Bonferroni-corrected significance thresholds were set to *P-value* < 1.1×10^-10^ (0.05/365,576/1,322) for plasma and *P*-value < 2.2×10^-10^ (0.05/365,576/628) for serum.

### Identification of independent association signals

To identify independent signals for metabolites, we performed conditional and joint association analyses using GCTA-COJO (v1.95.1)^34^ with the option of “--cojo-slct”. Genotypes from all individuals within either plasma or serum were used to construct the linkage disequilibrium (LD) reference panel for joint SNP effect estimation, which removed the effect shared among correlated SNPs due to LD between SNPs. The conditional significance threshold was set at 1.1×10^-10^ for plasma and 2.2×10^-10^ for serum, consistent with the primary mGWAS. Only the genomic window was specified to 5,000 kb with the parameter of “--cojo-wind 5000”, and all other parameters were set to their default values.

### Pairwise colocalization analysis of plasma and serum mQTL

To assess whether association signals for overlapping metabolites measured in plasma and serum were shared, we performed pairwise colocalization analysis between mQTL. When two mQTL for the same metabolite in plasma and serum were located within 150 kb of each other and showed strong LD (*r*^2^ > 0.8), colocalization analysis was conducted within the merged region (±500 kb around the mQTL) using the coloc.abf function in the coloc package (v5.2.3)^109^. The coloc package provides posterior probabilities for five mutually exclusive hypotheses regarding the associations of variants with two traits: H0: no association with either trait; H0: no association with either metabolite in plasma or in serum; H1: association with metabolite in plasma only; H2: association with metabolite in serum only; H3: association with same metabolite in plasma and serum due to linkage of two distinct causal variants; H4: association with same metabolite in plasma and serum due to a common single putative causal variant. A posterior probability for H4 (PP.H4) > 0.8 was applied to define significant colocalization between same metabolite in plasma and serum.

### Statistical fine-mapping

To delineate loci for statistical fine-mapping, we first defined mQTL regions as ±500 kb around each mQTL. Overlapping mQTL regions for the same metabolite were merged, and the resulting non-overlapping regions were subjected to fine-mapping analysis. We applied the Sum of Single Effects (SuSiE) regression framework to prioritize putative causal variants within each merged region using the susie_rss function^110^ implemented in the susieR package (v0.14.2)^111^. According to the susieR guidelines, the pairwise LD matrix for each region was calculated using PLINK based on genotype data from all individuals within the corresponding tissue. For each fine-mapping region, SuSiE provided posterior inclusion probabilities (PIPs) for each variant, representing the probability of being causally associated with the metabolite, as well as 95% credible sets (CSs) of putative causal variants constructed until the cumulative posterior probability exceeded 0.95. Variants with PIP > 0.5 were considered as putative causal variants.

### Stage-based mGWAS and mQTL analysis

To investigate stage-specific genetic effects on metabolite levels, we conducted stage-based mGWAS within either plasma or serum. The analysis framework was consistent with that described in the ***Genome-wide association study of plasma and serum metabolites***. For parity-based analysis, mGWAS was performed separately within each parity stage, and the parity term was removed from the model. Similarly, for lactation- based analysis, mGWAS was performed separately within each lactation stage, and the lactation term was removed from the model. To ensure adequate statistical power for mQTL detection, stage groups with fewer than 100 individuals were excluded. In plasma, three parity stages (heifer, primiparous, and multiparous) and four lactation stages (perinatal, early, peak, and end) were retained for analysis. In serum, two parity stages (primiparous and multiparous) and three lactation stages (early, peak, and end) were analyzed. Conditional analysis was subsequently performed within each stage group using the same procedures described in the ***Identification of independent association signals***.

### Context-sharing patterns of mQTL

To characterize the shared and specific architectures of mQTL across different contexts, we performed a meta-analysis of mQTL using the MashR package (v0.2.79)^35^. The analysis was restricted to mQTL identified by GCTA-COJO, and Z-scores (beta/se) derived from these signals were used as input. From the “mash” model, we obtained the posterior effect estimates and the corresponding significance levels, quantified as local false sign rate (LFSR). An mQTL was considered active in a given matrix or stage if LFSR < 0.05. To quantify pairwise similarity in the genetic regulation of metabolites across tissues or stages, we calculated Spearman’s correlation coefficients between posterior effect estimates, focusing on mQTL with LFSR < 0.05 in at least one tissue or stage.

### Integrative analysis of *cis*-eQTL and mQTL

To understand the potential molecular regulation between gene expression and metabolite levels, we integrated mQTL results with *cis*-eQTL data from the CattleGTEx resource. From the CattleGTEx Phase 1 database^32^, we selected *cis*-eQTL datasets from tissues with a sample size > 100, resulting in 29 tissues retained for downstream integrative analyses.

### Colocalization analysis

To identify shared putative causal variants between gene and metabolite, we conducted a colocalization analysis using the coloc.abf function implemented in the coloc package. For each metabolite, we defined mQTL regions as ±500 kb around each mQTL with the merged overlapping regions described in the ***Statistical fine-mapping***. Colocalization analysis were conducted between mQTL within the corresponding mQTL region and *cis*-eQTL for a given eGene. A PP.H4 > 0.8 was applied to define significant colocalization.

### Summary data-based Mendelian randomization (SMR) analysis

We performed summary data-based Mendelian randomization (SMR) analysis using SMR (v1.4.0)^40^ to evaluate putative causal or pleiotropic relationships between gene expression and metabolite level. Two models were tested: (i) E2M_SMR, in which gene expression were treated as the exposure and metabolite level as the outcome; and (ii) M2E_SMR, in which metabolite level were considered as the exposure and gene expression as the outcome. To prepare the input data, each mQTL region defined in the colocalization analysis was treated as a metabolite signal, and summary statistics for variants within each signal were converted to BESD format following the SMR data management. For *cis*-eQTL, only eGenes located within mQTL regions were tested, and the corresponding *cis*-eQTL summary statistics were likewise converted to BESD format. Only molecular QTL (*cis*-eQTL or mQTL) with a top nominal *P*-value < 1 × 10^-5^ were selected in the SMR test. Gene-metabolite pairs with *P*-value < 5 × 10^-6^ and HEIDI test *P*-value > 0.05 were selected and deemed as significant.

### Cell-type enrichment analysis

The Scpagwas software (v.1.3.0)^41^ was employed to perform enrichment analysis of cell type-metabolism associations. Briefly, scPagwas uses a polygenic regression model to prioritize a set of metabolism-relevant genes and uncover metabolism-relevant cell subpopulations by incorporating pathway-activity-transformed single-cell RNA-seq (scRNA-seq) data with mQTL summary statistics. In addition, 319 human KEGG pathways were retrieved from the KEGG database after eliminating duplicates and orthologous genes were converted to cattle orthologues. The Boot_evaluate function was employed to identify the significant metabolism-relevant cell types and calculate metabolism-relevant scores. The scGet_PCC function was used to prioritize the top metabolism-relevant genes based on ranked Pearson correlation coefficient (PCC). Genes ranked within the top 50 PCC values were defined as metabolism-relevant genes in each cell type. In addition, the scPagwas_perform_score function was applied to perform pathway activity analysis.

### Rare and low-frequency variant analysis of plasma metabolites

To assess the contribution of rare and low-frequency variants to plasma metabolite variation, we analyzed 2,910 individuals with plasma metabolites with LCWGS or HCWGS data. Variants with MAC < 3 and Hardy-Weinberg *P*-value < 1 × 10^-6^ were further excluded, leaving 16,834,909 high-quality SNPs for downstream analyses. Variants were classified according to MAF into rare variants (MAF < 0.005), low-frequency variants (0.005 ≤ MAF < 0.05) and common variants (MAF ≥ 0.05). We compared SNP heritability estimated using all variants with that estimated using common variants alone. For metabolites with significant SNP heritability based on all variants, the relative contribution of each MAF class was calculated as the proportion of total SNP heritability explained by the corresponding variant class. Single-variant association analyses were subsequently performed for plasma metabolites using all variants, following the same procedures as described in the ***Genome-wide association study of plasma and serum metabolites***.

Given the limited power of single-variant tests for rare and low-frequency variants, we further performed gene-based association analyses. Rare and low-frequency variant associations were tested using ACAT-V^49^ in GCTA software, which combines variant-level association *P*-values within each protein-coding gene using the aggregated Cauchy association test. All rare and low-frequency variants located within the boundaries of each protein-coding gene were included in the corresponding gene-based test. Common-variant gene-based associations were tested using MAGMA (v1.10)^50^ based on variant-level mGWAS summary statistics. In total, 19,462 protein-coding genes were tested for common variants and 19,970 protein-coding genes were tested for rare and low-frequency variants. Gene-metabolite associations were considered significant using Bonferroni-corrected thresholds of *P*-value < 0.05/19,462 for common-variant analyses and *P*-value < 0.05/19,970 for rare and low-frequency variant analyses.

### GWAS summary statistics of complex traits

To elucidate the regulatory mechanisms underlying complex traits in cattle, we integrated mQTL data with GWAS summary statistics from 11 complex traits in Holstein cattle, including five milk production traits, three reproduction traits, and three health traits (**Supplementary Table** 21). Independent SNPs were identified for each trait using GCTA-COJO, based on the corresponding LD reference panel.

### Two-sample Mendelian randomization

Two-sample Mendelian randomization (MR) analysis was performed to evaluate the potential causal effects of metabolites on complex traits. SNPs associated with the exposure (metabolites) and outcome (traits) were harmonized using the “harmonise_data” function in the TwoSampleMR package (v0.6.22)^55^. To obtain independent genetic variants associated with metabolites as instrumental variables (IVs) for MR analysis, we performed LD clumping on the mGWAS summary statistics using PLINK with the parameters of: clump-kb 500, clump-r2 0.1, clump-p1 1×10^-5^. MR analysis was conducted using the “mr” function in the TwoSampleMR package, applying four methods (inverse variance weighted, MR-Egger, weighted median, and weighted mode). In addition, heterogeneity was assessed using the MR-Egger method in the “mr_heterogeneity” function. Horizontal pleiotropy was evaluated by testing the MR-Egger intercept using the “mr_pleiotropy_test” function. Significant associations were defined as those with FDR-adjusted (*P*-value < 0.05) in at least two of the four MR methods, together with (*P*-value > 0.05) in both heterogeneity and horizontal pleiotropy tests.

### Enrichment analysis of GWAS signals

We used SumGSE^112^ to test whether GWAS signals were enriched in candidate regulatory regions relative to background regions across 11 complex traits. For mQTL, candidate regions were defined as the mQTL regions used in the ***Statistical fine-mapping analysis***. For *cis*-eQTL, candidate regions were defined as ±500 kb around lead *cis*-eQTL of each Gene, considering Whole Blood only. We then tested for enrichment of GWAS signals within eQTL and mQTL regions.

### Multi-trait colocalization analysis across eQTL, mQTL, and GWAS

We performed multi-trait colocalization analysis using the moloc package (v0.1.0)^113^ to identify shared putative causal variants among *cis*-eQTL, mQTL, and GWAS signals. Analysis were restricted to GWAS QTL regions overlapping mQTL regions. Prior probabilities were set to 1 × 10^-4^, 1 × 10^-5^, and 1 × 10^-6^ for the association of one, two, or three traits, respectively. For each region, moloc evaluated the evidence supporting 15 possible SNP-sharing configurations across the three traits. We considered the posterior probability that a shared putative causal variant was associated with all three traits (PPA.abc) > 0.5 as evidence for multi-trait colocalization.

### Comparative analysis of regulatory effects on metabolites between cattle and humans

To assess cross-species conservation of metabolite genetic regulation between cattle and humans, we collected publicly available human mGWAS summary statistics for plasma and serum metabolites. Plasma mGWAS summary statistics were obtained from the Canadian Longitudinal Study on Aging (CLSA) cohort, whereas serum mGWAS summary statistics were obtained from a meta-analysis of 2,901 Han Chinese individuals across three cohorts: the Zhejiang Metabolic Syndrome Cohort (ZMSC), the Westlake Precision Birth Cohort (WeBirth), and the Tongji-Huaxi-Shuangliu Birth Cohort (THSBC). Overlapping metabolites to those profiled in our study were matched using HMDB IDs or metabolite names, resulting in 37 plasma metabolites (**Supplementary Table** 26) and 38 serum metabolites (**Supplementary Table** 27).

The genomic coordinates of all human mGWAS variants were converted to the cattle reference genome (ARS-UCD1.2) using UCSC LiftOver (https://genome.ucsc.edu/cgi-bin/hgLiftOver). To quantify the proportion of heritability explained attributable to cattle loci orthologous to significant human mGWAS loci, we applied a two-random-effect LMM implemented in GCTA. For each metabolite, one GRM was specifically constructed using SNPs located within ±50 kb of cattle loci orthologous to significant human mGWAS loci (*P*-value < 1 × 10^-5^; referred to as human mQTL orthologous loci), where a second GRM was constructed using all remaining SNPs. The proportion of SNP heritability explained by human mQTL orthologous loci was subsequently compared with that explained by MAF-matched random loci.

In additional, we performed S-LDSC using LDSC (v1.0.1) to assess the contribution of human orthologs of cattle mQTL to human complex trait variation. GWAS summary statistics for 56 published human traits and diseases covering aging, metabolism, immune response, and female reproductive health were collected (**Supplementary Table** 28). Fine-mapped cattle plasma and serum mQTL variants were mapped to the human reference genome (hg38) using LiftOver. For each metabolite or predefined mQTL group, binary SNP annotations were constructed by marking human SNPs overlapping the orthologous cattle mQTL loci as annotated SNPs. S-LDSC was then used to estimate the proportion of SNP heritability explained by each annotation and to test whether cattle mQTL-derived annotations were enriched for heritability relative to the genome-wide SNP background. For stage-dependent analyses, mQTL were grouped according to parity or lactation stage before LiftOver and annotation construction, enabling enrichment tests for orthologous cattle mQTL active in specific physiological contexts. Multiple testing was controlled separately for plasma and serum annotations using Bonferroni correction.

## Data availability

The raw WGS and metabolome data newly generated in this study can be retrieved from GSA (https://ngdc.cncb.ac.cn/gsa/) and OMIX (https://ngdc.cncb.ac.cn/omix/). Accession number for WGS: CRA025735 (https://ngdc.cncb.ac.cn/gsa/s/fx09Xzs8); accession number for plasma metabolome: OMIX017044 (https://ngdc.cncb.ac.cn/omix/preview/Ej1Q8TTh); accession number for serum metabolome: OMIX017035 (https://ngdc.cncb.ac.cn/omix/preview/Fckh1Ff8). All mGWAS summary statistics were available to download at https://cattlema.farmgtex.org/download.

## Code availability

Code for the main analyses is freely available via GitHub at https://github.com/Tengjun0520/CattleMA_Pipeline.

## Acknowledgements

This work was supported by the National Key Research and Development Program of China (2021YFF1000701-06, 2023ZD0404901-05, and 2021YFD1200903-06), Taishan Scholar Foundation of Shandong Province (tsqnz20231240), Shandong Provincial Natural Science Foundation (ZR2023QC252), Agricultural Scientific and Technological Innovation Project of Shandong Academy of Agricultural Sciences (CXGC2025F09, CXGC2026C06, and CXGC2026G06), Shandong Cattle Research System (SDAIT-09-02), Key Research and Development Program of Shandong Province (2023LZGC004), and the Earmarked Fund for CARS (CARS-36). The funding bodies played no role in the design of the study and collection, analysis, and interpretation of data and in writing the manuscript. We thank the support of the SolAriot high-performance computing platform of the National Research Facility for Phenotypic and Genotypic Analysis of Model Animals (Beijing).

## Authors’ contributions

X. Wang and L. Fang conceptualized the study. L. Fang, X. Wang, J. Li, and Q. Zhang designed the experiments. J. Teng conducted the major data analysis. H. Li conducted the data analysis of cell type interaction part. J. Teng, H. Li, and X. Wang wrote the original manuscript. L. Fang substantially reedited and improved the manuscript. J. Yang, C. Duan, Z. Chen, and X. Zhang helped with data collection and integration. J. Lu contributed to web portal visualization. X. Zhao, F. Pei, X. Wu, P. Zhao, H. Zhang, and H. Gao contributed to data collection, experiment design and results interpretation. L. Guo, D. Wang, C. Ning, H. Liu, G. Su, R. Li, Y. Gao, J. Li, and Q. Zhang reviewed and improved the manuscript. L. Fang, X. Wang, and J. Teng edited the final version of the manuscript. All authors read and approved the final version of the manuscript.

